# A non-canonical function of LDHB promotes SLC7A11-mediated glutathione metabolism and protects against glutaminolysis-dependent ferroptosis in *KRAS*-driven lung cancer

**DOI:** 10.1101/2023.02.12.525859

**Authors:** Liang Zhao, Haibin Deng, Jingyi Zhang, Nicola Zamboni, Gerrit Adriaan Geest, Haitang Yang, Zhang Yang, Yanyun Gao, Duo Xu, Haiqing Zhong, Remy Bruggmann, Qinghua Zhou, Ralph A. Schmid, Thomas M. Marti, Patrick Dorn, Ren-Wang Peng

**Affiliations:** Division of General Thoracic Surgery, Inselspital, Bern University Hospital, University of Bern, Bern, Switzerland; Department for BioMedical Research (DBMR), University of Bern, Bern, Switzerland; Department of Biology, Institute of Molecular Systems Biology, Swiss Federal Institute of Technology/ETH Zürich, Zurich, Switzerland; PHRT Swiss Multi-Omics Center, smoc.ethz.ch, Zurich, Switzerland; Interfaculty Bioinformatics Unit and Swiss Institute of Bioinformatics, University of Bern, Bern, Switzerland; Lung Cancer Center, West China Hospital, Sichuan University, Chengdu, China; Department of Thoracic Surgery, Shanghai Chest Hospital, Shanghai Jiao Tong University, Shanghai, China; Department of Thoracic surgery, Fujian Medical University Union Hospital, Fuzhou City, Fujian, China; Department of Oncology, The First Affiliated Hospital of Nanjing Medical University, Nanjing, China

**Keywords:** Lactate dehydrogenase B (LDHB), Ferroptosis, Glutaminolysis, GPX4, SLC7A11, *KRAS*-dependent cancer

## Abstract

Ferroptosis, a form of non-apoptotic cell death program driven by excessive lipid peroxidation and an important mechanism of tumor suppression, is frequently dysregulated in cancer. However, the mechanisms underlying impaired ferroptosis in oncogene-specific tumors remain poorly understood. Here we report a non-canonical role of lactate dehydrogenase B (LDHB), whose main activity is the conversion of lactate to pyruvate, in protecting KRAS-mutated lung cancer from ferroptosis. Silencing of LDHB impairs intracellular glutathione (GSH) metabolism and drives the hypersensitivity of *KRAS*-mutant cells to ferroptosis inducers by inhibiting the SLC7A11/GSH/GPX4 axis, a central antioxidant system against lipid peroxidation and ferroptosis by catalyzing GSH synthesis and utilization. Mechanistically, LDHB promotes SLC7A11 expression and GSH biosynthesis, and inhibition of LDHB confers metabolic synthetic lethality with ferroptosis inducers due to increased glutaminolysis and production of reactive oxygen species (ROS) in mitochondria, ultimately triggering ferroptosis of *KRAS*-driven lung cancer cells. Consequently, combined inhibition of LDHB and SLC7A11 synergistically suppresses tumor growth in multiple *KRAS*-mutant lung cancer implants and in an autochthonous model of *Kras*-induced lung adenocarcinoma. Taken together, our results reveal a hitherto unrecognized mechanism of ferroptosis defense by glycolytic LDHB and suggest a new strategy for the treatment of *KRAS*-dependent lung cancer.

## INTRODUCTION

Oncogenic *KRAS* mutations are the most common genetic alterations in non-small cell lung cancer (NSCLC), particularly lung adenocarcinoma (LUAD) [1]. Despite advances in targeting the KRAS protein directly or indirectly [2, 3] and the advent of immunotherapy [4], effective therapies for *KRAS*-mutant lung cancer remain elusive [5, 6], arguing for a better understanding of *KRAS*-driven tumorigenesis and the development of novel therapeutic strategies to combat the disease.

KRAS mutations reprogram cancer metabolism, resulting in increased glucose uptake and glycolysis [7], altered glutamine/glutamate metabolism [8, 9], abnormal mitochondrial activity and redox equilibrium [10–12], which are critical for the increased energetic, biosynthetic, and redox requirements of tumor cells and therefore required for KRAS-induced tumorigenicity [13, 14]. In particular, *KRAS*-mutant cancer produces elevated levels of reactive oxygen species (ROS) and has evolved complex antioxidant programs to overcome the oxidative stress barrier during tumorigenesis [15, 16], on which tumor cells strongly depend for survival [10-12, 17, 18]. Consequently, disruption of ROS defense would be selectively toxic for cancer cells [18–21]. Although it is generally accepted that metabolic ROS play a key role in the development and progression of *KRAS*-driven cancers, the mechanism by which ROS are scavenged or eliminated are not fully understood [11].

Lactate dehydrogenase B (LDHB or LDH1) is a subunit of LDH, a tetrameric enzyme consisting of LDHB and LDHA (LDH2). While LDHA catalyzes the reduction of pyruvate to lactate, LDHB preferentially oxidizes lactate to pyruvate, coupled with NAD^+^ to NADH [22, 23]. Despite different functions, LDHA and LDHB are important for ATP generation and homeostasis under physiologically low O2 (anaerobic glycolysis) and also for the reprogramed cancer metabolic state of aerobic glycolysis known as the Warburg effect [24, 25]. LDHB has been shown to be essential for *KRAS*-mutant lung cancer progression [25–27] and to regulate stem cell properties and mitochondrial functions [28]. LDHB may have other functions independent of its role in lactate metabolism [23, 29, 30]. However, the mechanisms by which LDHB contributes to *KRAS*-mutant NSCLC is largely unknown.

Ferroptosis, a non-apoptotic regulated cell death program activated by ROS- and iron-dependent lipid peroxidation of polyunsaturated fatty acid (PUFA) [31, 32], is often dysregulated in cancer [33]. The susceptibility of cancer cells to ferroptotic death is determined by the antagonism between the cellular metabolic activities that trigger lipid oxidation and the cellular antioxidant systems that counteract it [34]. The solute carrier family 7 member 11 (SLC7A11; also known as xCT) and the selenium-dependent hydroperoxidase glutathione peroxidase 4 (GPX4) are the most potent antioxidant hubs protecting against ferroptosis [35, 36], with the cystine/glutamate antiporter subunit SLC7A11 importing cysteine to synthesize glutathione (GSH), a critical cofactor to multiple antioxidant proteins, including GPX4, thereby detoxifying lipid peroxides and suppressing ferroptosis [33, 37]. Blocking this antioxidant axis with SLC7A11 and GPX4 inhibitors (ferroptosis inducers) such as the small-molecule Erastin and RSL3, respectively, leads to uncontrolled accumulation of lipid peroxides at the plasma membrane and other endomembranes within the cell, which promotes membrane permeabilization, damages membrane integrity and ultimately induces ferroptosis [34, 38]. *KRAS*-mutant cancers exhibit a pronounced dependence on antioxidant adaptations [39, 40], and evasion or inactivation of ferroptosis contributes to Kras-driven tumor development and progression [41, 42]. However, the cellular processes and context-specific factors that modulate ferroptosis sensitivity of *KRAS*-dependent lung cancer cells by impinging on key antioxidant proteins such as SLC7A11 and GPX4 are poorly understood.

In this study, we report the unexpected finding that LDHB protects against ferroptosis in *KRAS*-mutant lung cancer. LDHB promotes SLC7A11 expression and GSH metabolism, implicating a role independent of its canonical function in lactate metabolism. Specifically, LDHB depletion impairs GSH synthesis and leads to hypersensitivity of *KRAS*-mutant cancer cells to ferroptosis inducers, which is causally linked to increased glutaminolysis and mitochondrial ROS. Combined inhibition of LDHB and SLC7A11 synergistically suppresses the growth of *KRAS*-mutant cancer cells and tumors by inducing ferroptotic death *in vitro* and *in vivo*, including a genetically engineered mouse model of Kras-induced LUAD with Ldhb deficiency. Our results uncover a previously unrecognized mechanism of ferroptosis defense in *KRAS*-mutant lung cancer and suggest a new strategy for the treatment of the disease.

## RESULTS

### LDHB regulates glutathione metabolism in *KRAS*-dependent lung cancer cells

To identify metabolic pathways that underpin the function of LDHB in *KRAS*-dependent lung cancer [27, 28], we performed unbiased metabolomics of LUAD cells carrying *KRAS* mutations (A549, H358). Knockdown (KD) of LDHB by small-interfering RNAs (siRNAs) significantly altered numerous cellular metabolites (**Figure 1a; Supplementary Figure 1a-f**), which were enriched in multiple metabolic processes, among which amino acid metabolisms (e.g., glutamate metabolism, glycine and serine metabolism, and cysteine and methionine metabolism) being the most significantly downregulated pathways upon LDHB KD (**Figure 1b**). In particular, key amino acids or intermediate metabolites involved in *de novo* or salvage synthesis of glutathione (GSH), particularly cysteine (Cys), γ- glutamylcystine (γ-GC), glutamine (Gln), glutamate (Glu), and GSH itself, were significantly decreased in LDHB KD A549 cells compared with those in control A549 cells expressing control siRNA (siNT) (**Figure 1c, d**). Similar results were observed in H358 cells (**Figure 1e, g; Supplementary Figure 1a-f; Supplementary Table 1**).

**Figure 1.**
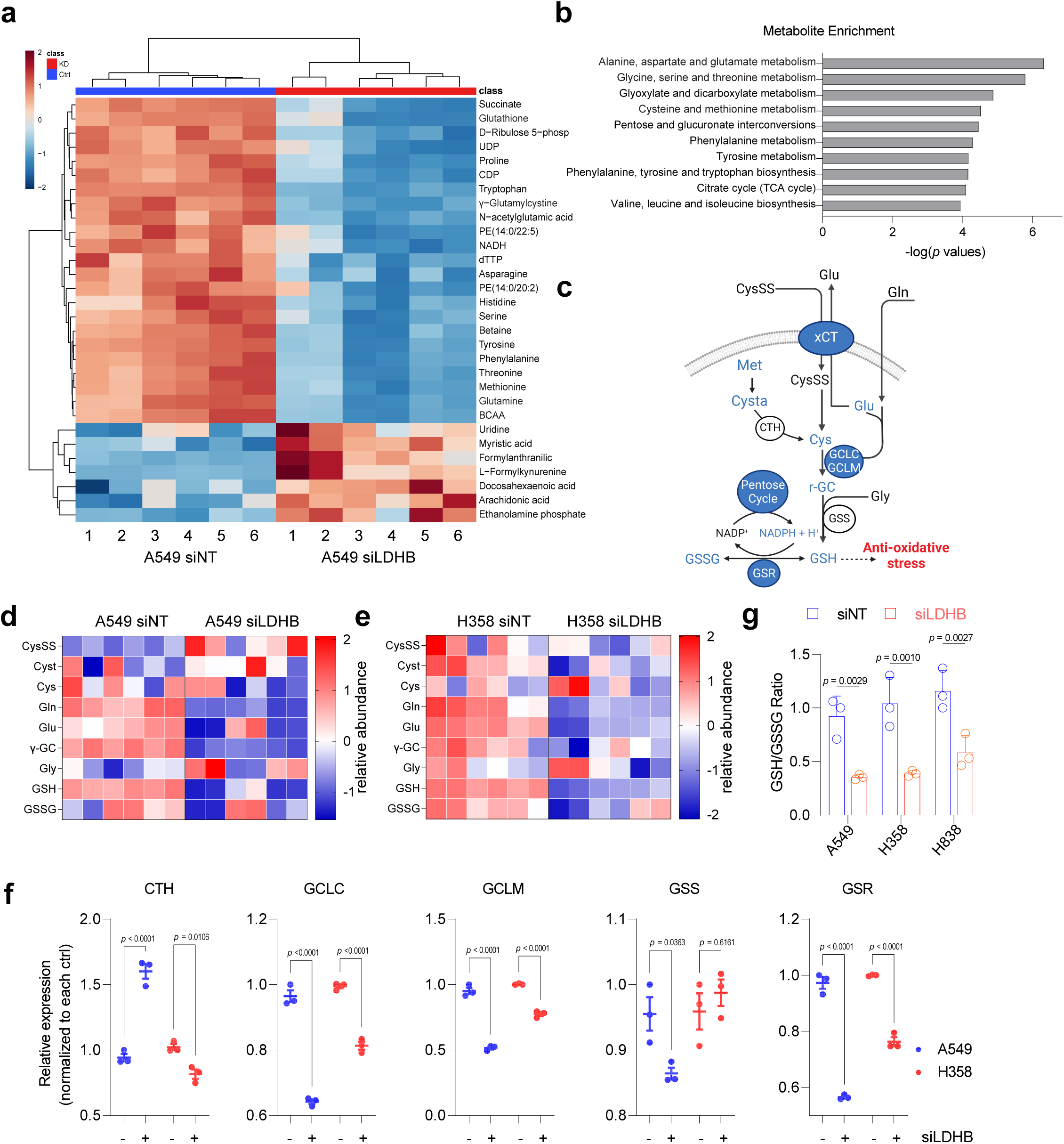
LDHB silencing impairs GSH metabolism in *KRAS*-dependent lung cancer cells. **a.** Heat map showing the top 30 significantly different metabolites between A549 (siNT) and A549 (siLDHB) cells (n=12). Metabolomics analysis was performed 48 hours after siRNA transfection. Relative abundance is scaled between 2 to -2. **b.** Pathway enrichment analysis shows significantly downregulated metabolic processes in LDKB KD A549 cells (siLDHB) compared to control A549 cells (siNT). **c.** Schematic showing the regulatory effect of LDHB on de novo GSH synthesis. Enzymes, ion channels, and metabolic pathways (e.g., PPP) are marked in circles, among which the expression of corresponding genes and metabolites that are significantly altered by LDHB KD in both A549 and H358 cells are highlighted in blue. **d-e,** Heat map illustrating the abundance of key metabolites contributing to GSH synthesis. **f**, The fold change of mRNA levels of key GSH synthesis enzymes (CTH, GCLC, GCLM, GSS and GSR) after LDHB KD. CysSS, cysteine; Glu, glutamate; Gln, glutamine; XCT, SLC7A11/SLC3A2; Cys, cysteine; Met, methionine; Cysta, cystathionine; CTH, gamma- cystathionase; GSR, glutathione; GSSG, glutathione disulfide; GCL, glutamate-cysteine ligase; γ-GC, γ-glutamylcysteine; Gly, glycine; GS, glutamine synthetase; GSR, glutathione reductase. The *p* values are calculated by two-way ANOVA. **g,** LDHB KD impairs GSH/GSSG ratios in A549 and H358 cells. Intracellular GSH and GSSG levels were measured 48 hours post transfection with siRNAs (siNT and siLDHB). Data are presented as mean ± s.d. (n=3), with *p* values by two-way ANOVA.

Interestingly, LDHB KD led to increased cystine (CysSS) but decreased GSH disulfide (GSSG; the oxidized form of GSH) in A549 and H358 cells, suggesting that LDHB may also regulate the conversion of cystine to cysteine, a metabolic step dependent on sufficiently levels of NADPH. LDHB is important for the Warburg effect [24, 25] and the pentose phosphate pathway (PPP), a branch of glucose metabolism that degrades glucose 6-phosphate (G6P) to ribose 5-phosphate (R5P) and to lactate[43], is an important resource for the production of NADPH. To monitor glycolysis [G6P - glucose 3- phosphate (G3P) - lactate] and PPP activities in more detail, we performed targeted metabolic profiling using ^13^C-labelled glucose. LDHB KD significantly decreased the PPP-derived lactate but increased glycolysis-derived lactate (**Supplementary Figure 2**), suggesting that LDHB KD reduces PPP activity, impairs NADPH synthesis and limits the conversion of cysteine to cysteine.

To corroborate the observations from the metabolomics analysis, we performed whole-genome transcriptomic profiling (RNA sequencing) of A549 and H358 cells. LDHB KD significantly downregulated GSH synthesis genes, such as glutamate-cysteine ligase (GCL) catalytic and modifier subunits (GCLC and GCLM), GSH synthetase (GSS), and GSH reductase (GSR), as well as the GSH gene signature (**Figure 1f; Supplementary Figure 3a-c**). Interestingly, cystathionine-γ-lyase (CTH), the key enzyme in *de novo* cysteine synthesis that generates cysteine from methionine-derived homocysteine and serine as a product of the trans-sulfuration pathway, was also reduced (**Figure 1f; Supplementary Figure 3b**), consistent with our metabolomics data that LDHB KD decreased glycine and methionine metabolism (**Supplementary Figure 1d, f**).

To further validate the metabolomics and transcriptomic results, we determined the ratio of reduced GSH to oxidized GSH (GSH/GSSG), a standard measure of cellular oxidative stress. LDHB KD significantly and unanimously decreased the GSH/GSSG ratio in A549, H358 and H838 cells, indicating that LDHB deficiency impairs GSH synthesis (**Figure 1g**). Taken together, these results suggest that LDHB regulates GSH metabolism in *KRAS*- dependent NSCLC cells.

### LDHB suppression drives hypersensitivity of KRAS-dependent lung cancer cells to ferroptosis inducers

Our finding that LDHB KD upregulates oxidative stress in *KRAS*-mutant lung cancer cells prompted us to identify the cellular processes that represent selective vulnerabilities of LDHB-depleted cells. Synthetic lethal chemical screens in a panel of lung cancer cell lines and a normal epithelial cell line (BEAS-2B) with small-molecule drugs (n=22) targeting various oncogenic and metabolic pathways (**Supplementary Table 2**) identified Erastin, an inhibitor of SLC7A11 and potent inducer of ferroptosis by blocking the uptake of cystine for GSH synthesis, that synergized with LDHB-specific siRNAs to suppress the viability of a subset of *KRAS*-mutant lung cancer cells (A549, H838, H460, H2122, and H2009), despite to a varied degree (**Figure 2a, b; Supplementary Figure 4e**). Clonogenic assay confirmed the synergistic effect of LDHB KD and Erastin in *KRAS*-dependent human lung cancer cells and murine KP (*Kras^G12D/+;^ p53^-/-^*) cells (**Figure 2c; Supplementary Figure 4a**).

**Figure 2.**
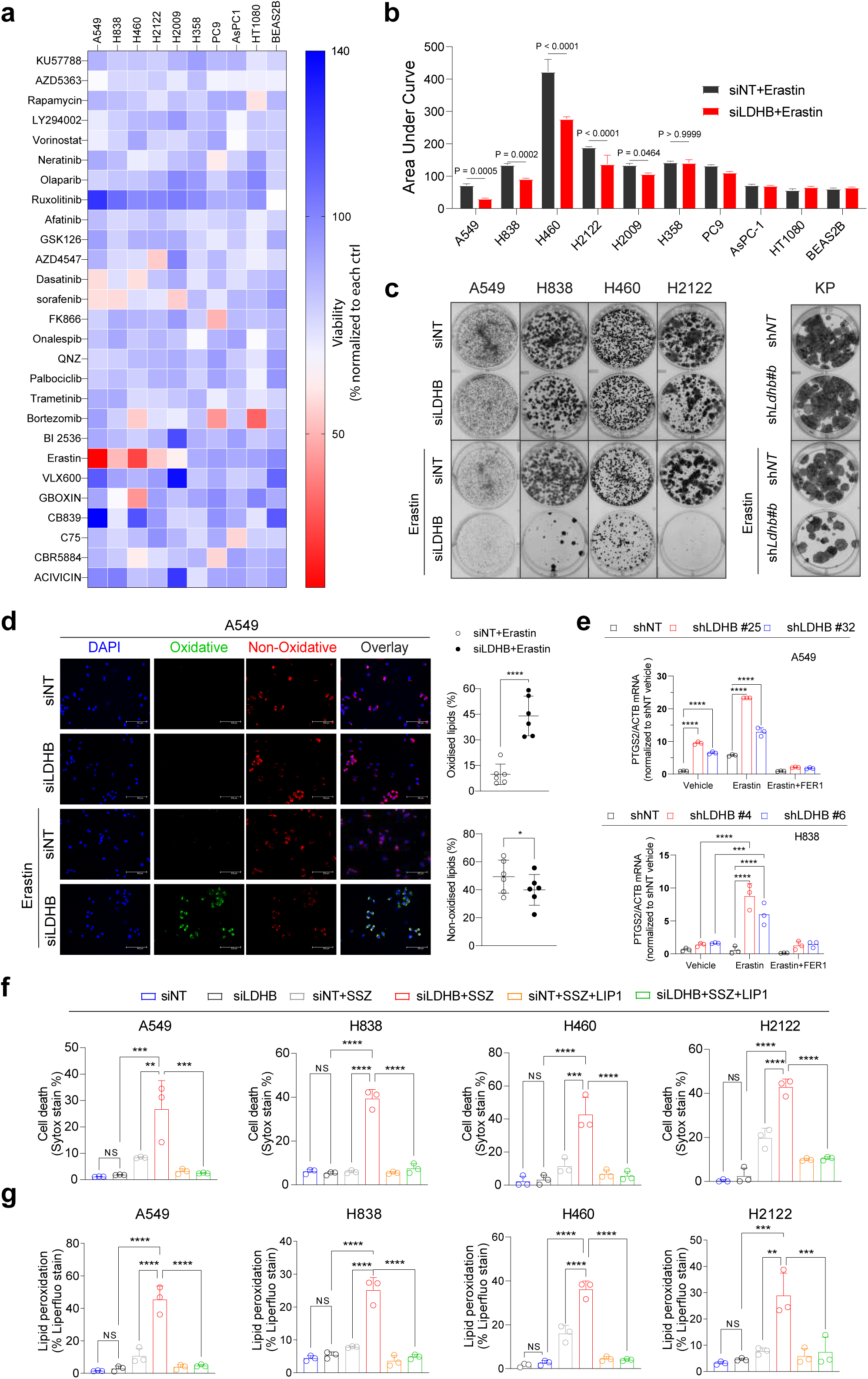
SLC7A11 is a synthetic lethal target in LDHB-deficient *KRAS*-mutant lung cancer cells. **a,** Heat map illustrating relative cell viability of LDHB KD cancer and normal epithelial cell lines after treated for 72 hours with the indicated anti-cancer and metabolic compounds. Data are presented as percentages of viable LDHB KD cells (expressing siLDHB) normalized to viable control cells that express siNT. **b,** Sensitivity of LDHB KD cells (siLDHB) and control cells (siNT) to the ferroptosis inducer erastin. Drug sensitivity is determined by the value of area under curve (AUC). Data are presented as mean ± s.d. (n=3), with p < 0.05 by two-way ANOVA with Tukey’s multiple comparison test. **c,** Lung cancer cell lines (A549, H838, H460, and H2122) and murine KP cells (*Kras^G12D/WT^*; *p53^fl/fl^*) were transfected by LDHB-targeted siRNAs (siLDHB) and control siRNAs (siNT) for 24 h, further treated for 72 h with DMSO or Erastin and subjected to clonogenic assay. **d,** A549 cells were transfected with siNT or siLDHB for 36 h, further treated with vehicle (DMSO) or Erastin (5 μM) for 14 h before stained with C11 BODIPY 581/591 and DAPI. Oxidized and non-oxidized lipids were analyzed by fluorescence microscopy. Scale bars, 100 μm. **e,** Quantitative analysis (RT-PCR) of *PTGS2* mRNA levels in the indicated cells after treatment for 14 h with vehicle (DMSO) or Erastin in the presence or absence of FER1 (ferroptosis inhibitor). **f, g,** Cell death (Sytox stain; f) and lipid peroxidation (liperfluo stain; g) assay of the indicated cancer cells treated with siNT, siLDHB for 48 hours, followed by further treatment for 16 hours with sulfasalazine (SSZ), alone or in the presence of the ferroptosis inhibitor Liproxstatin-1 (LIP1). Data are shown as mean ± s.d. (n=3). *p < 0.05. ** p < 0.01. ***p < 0.001. ****p < 0.0001 by one-way ANOVA.

Next, we investigated whether LDHB suppression promotes Erastin-induced ferroptosis. Lipid peroxidation (C11-BODIPY staining) assay showed that LDHB KD A549 cells had similar levels of lipid peroxidation and mitochondrial superoxides as control cells (**Figure 2d, 4f**; **Supplementary Figure 8f**), but Erastin significantly increased oxidized lipids but decreased non-oxidized lipids in LDHB KD A549 cells (**Figure 2d**). Consistently, the expression of PTGS2, a ferroptosis biomarker [19], was induced by LDHB KD in A549 and H838 cells, and Erastin further pronounced the increase (**Figure 2e**). Importantly, the upregulation of PTGS2 incurred by LDHB KD alone or its combination with Erastin was effectively abrogated by the ferroptosis inhibitor FER1 (**Figure 2e; Supplementary Figure 4h**), which eliminates lipid hydroperoxides and produces a similar anti-ferroptotic effect as GPX4[35, 38]. In addition, LDHB KD plus Erastin or RSL3 synergistically inhibited A549, H838, and H460 the viability, which could be rescued by FER1, partially by the necrosis inhibitor (NEC) due to the shared features of the two processes [44], but not by inhibitors of autophagy (HCQ) or apoptosis (ZVF) (**Supplementary Figure 4b, c, d**).

Similar results were observed with sulfasalazine (SSZ), an FDA-approved drug with ferroptosis-inducing activity. SSZ mildly induced cell death and lipid peroxidation in A549, H838, H460, and H2122 cells, but it caused a significantly greater percentage of cell death and lipid peroxidation in LDHB KD cells (**Figure 2f, 2g, Supplementary Figure 4k**). Again, the potent inhibitory effect of SSZ on cell death and lipid peroxidation in LDHB KD cells was effectively counteracted by the ferroptosis inhibitor LIP1 (**Figure 2f, g**). Similar results were obtained when ML162, an independent GPX4 inhibitor, was used (**Supplementary Figure 4i, j, k**). Thus, LDHB protects KRAS-dependent lung cancer cells from oxidative stress and LDHB suppression drives hypersensitivity to SLC7A11/GPX4 inhibition-induced ferroptosis.

### LDHB upregulates the expression of SLC7A11 through the INFγ-JAK-STAT1 axis

To elucidate the mechanisms by which LDHB regulates GSH synthesis and ferroptosis defense, we re-examined our RNAseq data of A549 and H838 cells [28]. LDHB KD affected not only GSH biosynthesis genes (**Figure 1g**), but also those modulating ferroptosis sensitivity, with the ferroptosis resistance gene signature significantly attenuated in LDHB KD A549 and H838 cells, among which SLC7A11 decreased the most (**Figure 3a, Supplementary Figure 6a**). Confirming the RNAseq data, LDHB KD markedly decreased SLC7A11 protein levels in A549, H838, H460, H2122, murine KP cells (*KRAS^G12D^*, *p53^−/−^*), and A549 xenograft tumors (**Figure 3b, c; Supplementary Figure 6b**).

**Figure 3.**
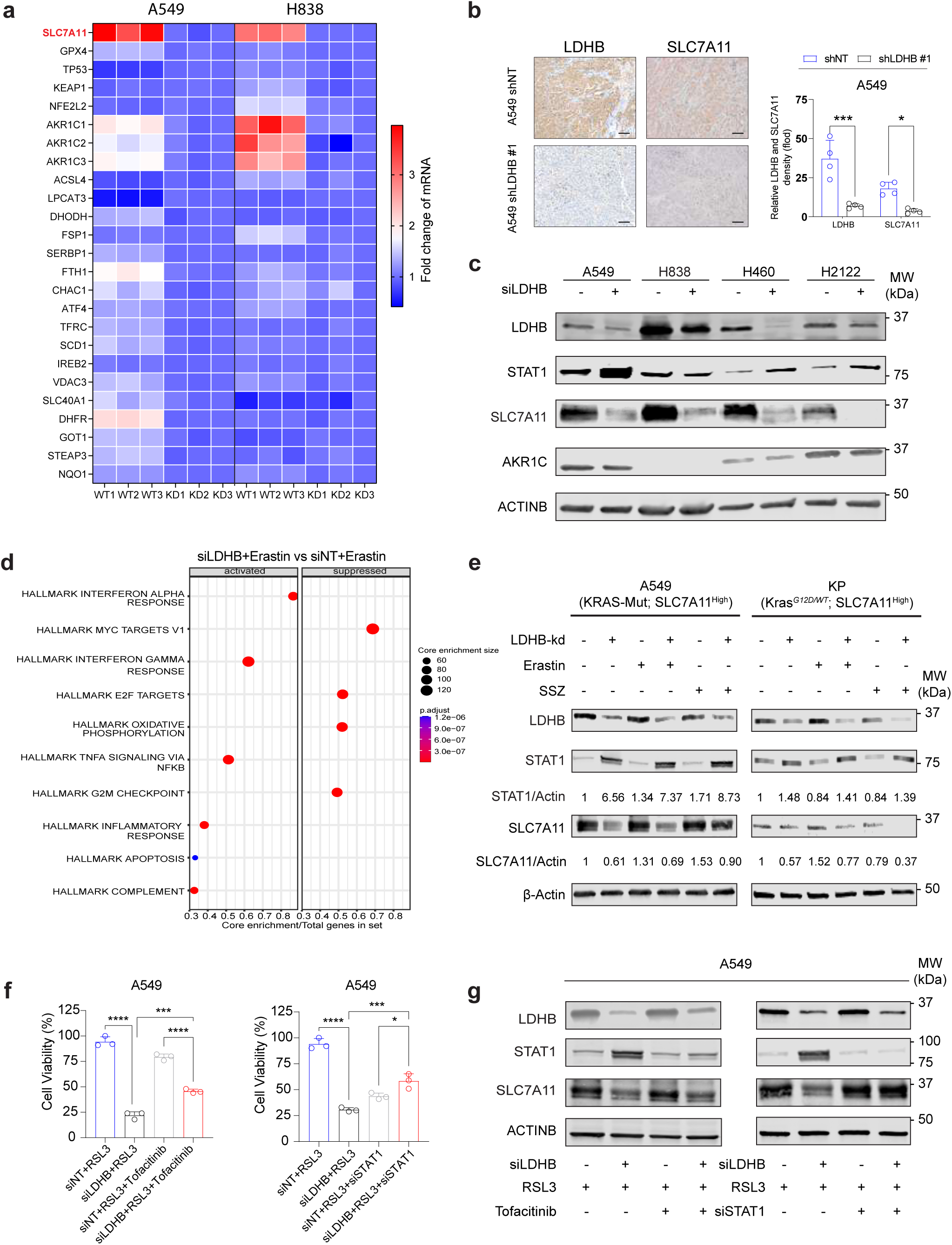
LDHB deficiency downregulates SLC7A11 expression and sensitizes *KRAS*-driven lung cancer cells to ferroptosis inducers. **a,** Heat maps showing the fold change (mRNA level) of ferroptosis genes (n=25) in siLDHB cells (KD) compared to siNT cells (WT). **b,** Immunohistochemical analysis of LDHB and SLC7A11 in xenograft tumors derived from LDHD KD A549 cells. Quantification is shown to the right, with \**p*<0.01, \*\*\**p*<0.001 by two-way ANOVA. **c,** Immunoblots of the indicated cancer cells transfected for 48 h with siNT (-) and siLDHB (+). **d,** Gene set enrichment analysis (GSEA) of the transcriptome of A549 cells treated with LDHB-specific siRNAs and Erastin (siLDHB+Erastin) compared to that of A549 cells treated with control siRNAs and Erastin (siNT+Erastin). **e,** Immunoblots of human A549 and murine KP cells transfected (48 h) with LDHB-specific siRNAs (KD) and control siRNAs (-) followed by further treatment for 16 hours with Erastin (5 uM) or Sulfasalazine (1 mM). **f,** Viability assay of A549 cells treated (24 h) with LDHB-specific siRNAs (siLDHB), STAT1-specific siRNAs (siSTAT1) and control siRNAs (siNT) followed by further treatment 24 hours with RSL3 or Tofacitinib, alone or in combination. Data are presented as mean ± s.d. (n=3), with P<0.01, ***P<0.001 by two-way ANOVA. **g,** Immunoblots of A549 cells transfected with LDHB-specific siRNAs (siLDHB) for 24 hours, then STAT1-specific siRNAs (siSTAT1) and control siRNAs (-) followed by further transfection for 48 hours; Or Immunoblots of A549 cells transfected with LDHB-specific siRNAs (siLDHB) for 48 hours, and then treated with RSL3 or Tofacitinib, alone or in combination for 24 hours.

To validate SLC7A11 as the target by which LDHB negatively regulates ferroptosis, we overexpressed SLC7A11 in A549 and H838 cells in which LDHB was stably knocked down by shRNAs (**Supplementary Figure 7a**). SLC7A11 overexpression increased the resistance to Erastin and SSZ and concomitantly decreased lipid peroxidation in LDHB KD cells (**Supplementary Figure 7b, c, d, e**). Consistently, N-acetyl cysteine (NAC), a widely used antioxidant in clinical studies, abrogated the combinatorial effect of LDHB KD and Erastin in A549 and H838 cells (**Supplementary Figure 7d, e**). These results suggest that LDHB depletion limits GSH biosynthesis and promotes ferroptosis by downregulating SLC7A11.

To investigate how LDHB regulates SLC7A11, we transcriptomically profiled A549 cells treated with scrambled siRNAs (siNT), LDHB-specific siRNAs (siLDHB), and Erastin, alone and in combination. We identified 312 genes that were decreased (log2FoldChange ≥ 2) by the combination (siLDHB/Erastin) compared to Erastin alone (siNT/Erastin), of which 296 were statistically significant (p-value < 0.05) (**Supplementary Figure 6c; Supplementary Table 3**). Pathway enrichment analysis revealed “interferon α/γ (INFα/γ) responses” as top candidate significantly upregulated/activated by Erastin in siLDHB A549 cells (**Figure 3d, Supplementary Figure 6d**). INFα/γ signaling plays a key role in immune response, and its activation is mediated by Janus kinase (JAK) and signal transducer and activator of transcription (STAT) [45]. The INFγ-JAK-STAT1 pathway has been shown to impair cystine uptake and promotes lipid peroxidation and ferroptosis through transcriptional repression of the system xc^-^ subunits SLC3A2 and SLC7A11 [45]. Therefore, we tested the possibility that LDHB regulates SLC7A11 via STAT1. LDHB KD markedly increased STAT1 protein levels in A549, H358, and murine KP cells, and this increase was accompanied by SLC7A11 decrease (**Figure 3e, Supplementary Figure 6b**). Remarkably, xCT-targeted drugs (Erastin and SSZ) only slightly affected SLC7A11 protein levels in A549 and KP cells but caused a greater decrease in LDHB KD cells (**Figure 3e, Supplementary Figure 6e**). In contrast, LDHB KD, alone or in combination with Erastin or SSZ, failed to apparently affect STAT3, SCD1, C-MYC, or acyl-CoA synthetase long-chain family member 4 (ACSL4), which have been reported to context-dependently regulate ferroptosis [46–49] (**Supplementary Figure 6f**).

Finally, we examined whether the INFγ-JAK-STAT1 pathway is causally linked to RSL3- induced ferroptosis in LDHB KD cells. Blocking the pathway by the JAK inhibitor tofacitinib or STAT1 KD abrogated the inhibitory effect of RSL3 on the viability of LDHB KD A549 cells (**Figure 3f**) and restored the protein level of SLC7A11 that was reduced by RSL3 in LDHB KD cells (**Figure 3g**).

Since both LDHA and LDHB are required for the Warburg effect [50] and LDHA has also been reported to play a role in NSCLC [51] and in ROS alleviation [52], we investigated whether LDHA has similar function as LDHB in ferroptosis defense. LDHA KD did not appear to affect STAT1 expression or the sensitivity to the ferroptosis inducers RSL3 and Erastin in A549 and H838 cells (**Supplementary Figure 5a, b, c**), as did the LDHA inhibitors (GSK2837808A and R-GNE-140) (**Supplementary Figure 5d, e**), suggesting that LDHA is not involved in ferroptosis surveillance in KRAS-mutant NSCLC.

Taken together, these results suggest that LDHB promotes ferroptosis evasion in *KRAS*- dependent lung cancer by upregulating the expression of SLC7A11, which is achieved through negative regulation of the INFγ-JAK-STAT1 axis.

### SLC7A11 inhibition-induced ferroptosis in LDHB-deficient lung cancer cells requires glutaminolysis

Next, we explored the metabolic process that underpins the ferroptosis provoked by LDHB/SLC7A11 inhibition. It has been shown that SLC7A11 antagonizes glutamine metabolism [53] and that glutaminolysis, a mitochondrial process that converts glutamine to metabolites to fuel the tricarboxylic acid (TCA) cycle, is required for the execution of ferroptosis [54]. As LDHB regulates SLC7A11, we posit that LDHB/SLC7A11 inhibition-induced ferroptosis activates glutaminolysis. Indeed, transcriptomic profiling indicated that SLC7A11 inhibition by Erastin significantly upregulated the expression of gluataminolysis genes in A549 cells and LDHB KD A549 cells (**Figure 4a, b**). In line with this finding, Erastin significantly reduced GSH-related metabolites (Cysteine, Cys-Gly, GSSG) but increased glutamine uptake in LDHB KD A549 cells (**Figure 4c, Supplementary Figure 8a, b,**), suggesting that combined inhibition of LDHB and SLC7A11 increased glutaminolysis.

**Figure 4.**
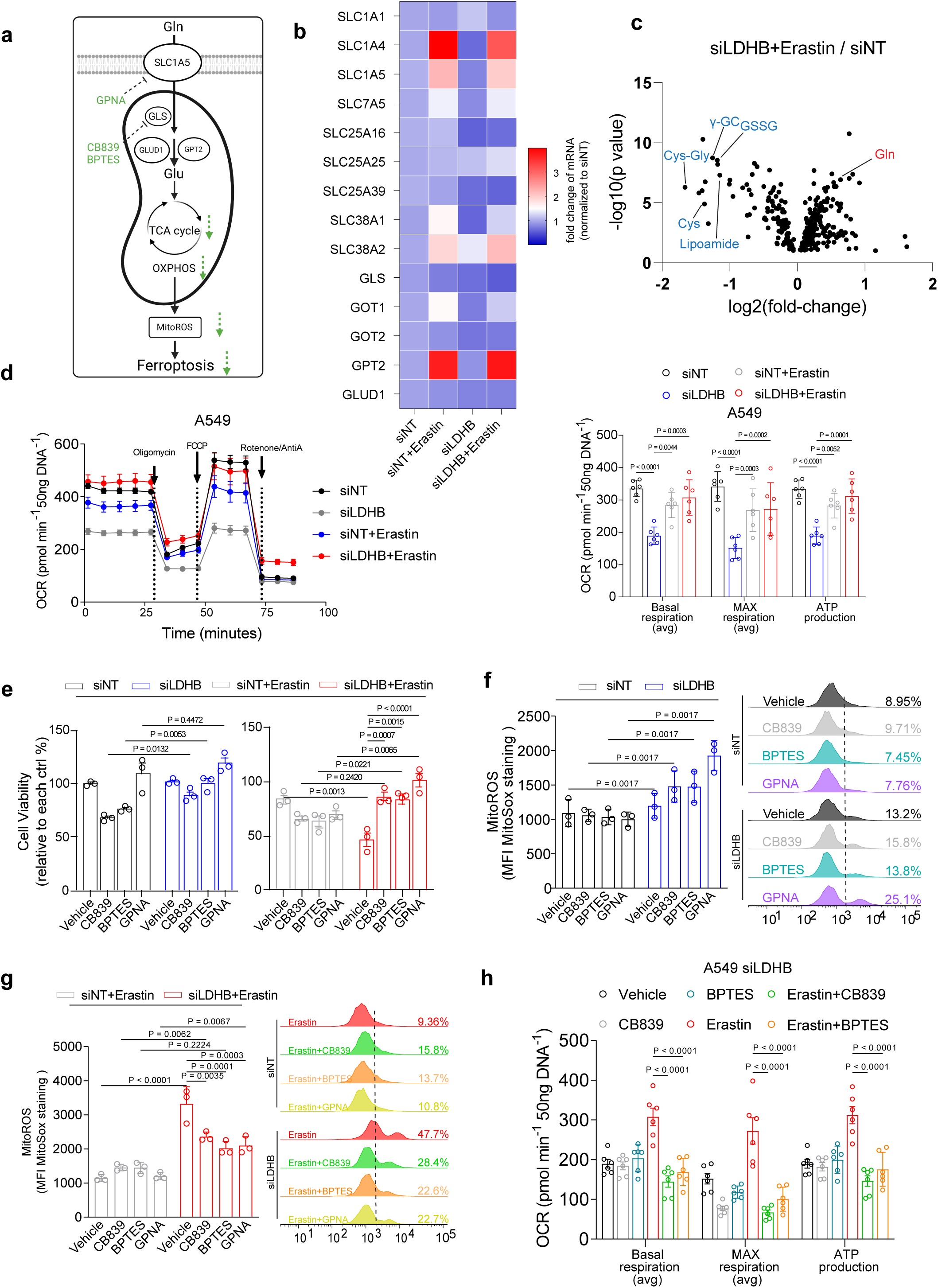
Glutaminolysis is required for the metabolic synthetic lethality between LDHB inactivation and ferroptosis inducer. **a,** Schematic of glutaminolysis and its contribution to LDHB/SLC7A11 inhibition-induced ferroptosis. Enzymes and glutamine transporters are marked in circles, and inhibitors that mitigate LDHB/SLC7A11 inhibition-induced ferroptosis are highlighted in green. **b,** Fold changes (mRNA levels) of glutaminolysis genes in A549 cells transfected for 48 hours with LDHB-specific siRNAs (siLDHB) or control siRNAs (siNT), followed by further treatment with Erastin and vehicle for 20 hours. **c,** Volcano plot comparing the metabolomics profile of A549 cells transfected for 48 hours with siNT or siLDHB and further treated with Erastin or vehicle for 20 h. **d,** Oxygen consumption rate (OCR) of A549 cells transfected and treated as above (b-c). Data are presented as mean ± s.e.m (n=6). **e,** Viability assay of A549 cells transfected with siNT or siLDHB for 48 hours and further treated for 20 h with DMSO, Erastin (5 μM), CB839 (0.5 μM), BPTES (2 μM), and GPNA (50 μM), alone or in combination. Normalization was based on vehicle-treated siNT and siLDHB groups (set as 100%), respectively. **f-g,** MitoSox (mitochondrial ROS marker) quantification (left) and flow cytometry (right) analysis of A549 cells transfected and treated as in b-e; Quantification of mitoROS are shown as mean fluorescence intensity ± s.d. (n=3), with siNT group used for normalization (left). **h**, Oxygen consumption rate (OCR) of A549 cells transfected and treated as in b-g. Data are shown as the mean ± s.d. (n=6), with statistical analyses by two-way ANOVA.

Glutaminolysis fuels the TCA cycle and subsequent oxidative phosphorylation (OXPHOS) in mitochondria, we therefore determined whether combined inhibition of LDHB/SLC7A11 leads to abnormal respiration and energy production in mitochondria. Using the Seahorse XF Mito Stress Analyzer, a real-time measure of the oxygen consumption rate (OCR), LDHB KD (siLDHB) indeed markedly reduced OCR and mitochondrial ATP in A549 cells, consistent with its role in mitochondrial metabolism [25, 28]. Remarkably, Erastin highly increased OCR in LDHB KD (siLDHB + Erastin) but not in control A549 cells (siNT + Erastin) (**Figure 4d, Supplementary Figure 8c**), suggesting that cancer cells subjected to combined inhibition of LDHB and SLC7A11 undergo a high level of OXPHOS.

To determine whether glutaminolysis directly contributes to LDHB/SLC7A11 inhibition-evoked ferroptosis, we investigated functional importance of several key glutaminolysis enzymes in this process. Suppression of glutaminolysis by inhibiting GLS (CB839; BPTES) or SLC1A5 (GPNA), the key enzyme and glutamine transporter whose activities are crucial for the process (**Figure 4a**), substantially rescued the viability of Erastin-treated LDHB KD A549 cells (**Figure 4e**), indicating that LDHB KD/Erastin inhibits cell viability through glutaminolysis. Notably, the viability increase upon glutaminolysis inhibitors (CB839, BPTES, and GPNA) was accompanied by a corresponding decrease in mitoROS, a measure of mitochondrial superoxides (**Figure 4f, g**). Concomitant with mitoROS change, both CB839 and BPTES significantly decreased OCR in LDHB KD cells treated with Erastin (**Figure 4h; Supplementary Figure 8d, e, f, g**). Thus, the combined inhibition of LDHB and SLC7A11 hyperactivates glutaminolysis, resulting in the accumulation of mitochondrial ROS and induction of ferroptosis. These results suggest that LDHB and SLC7A11 inhibition induces ferroptosis by increasing glutaminolysis-dependent ROS production in *KRAS*-mutant cancer cells.

### Combined inhibition of LDHB and SLC7A11 synergistically suppresses *KRAS*-mutant tumor growth by inducing ferroptosis *in vivo*

We validated our *in vitro* findings in *KRAS*-mutant lung cancer xenografts and a genetically engineered mouse (GEM) model of autochthonous *Kras^G12D^*-induced lung adenocarcinoma that accurately resembles human disease [20]. In implantable xenograft models (LDHB KD A549 and H838), the SLC7A11 inhibitor Erastin (administrated at 30mg/kg) significantly and consistently suppressed the growth of LDHB KD tumors, despite marginal effects on control A549 and H838 tumors (**Supplementary Figure 9a**). Similar results were observed with the FDA-approved SLC7A11 inhibitor SSZ (150 mg/kg) that substantially suppressed the growth of LDHB KD H460 tumors but not control H460 tumors (**Figure 5a**, **b**). In parallel with tumor shrinkage, Erastin significantly upregulated the levels of 4-HNE, a lipid peroxidation marker, but not cleaved caspase-3 and Ki67, in LDHB KD tumors, reinforcing the notion that ferroptosis underpins the anti-tumor effects of SLC7A11 inhibition in LDHB KD cancer cells (**Supplementary Figure 9c, e**).

**Figure 5.**
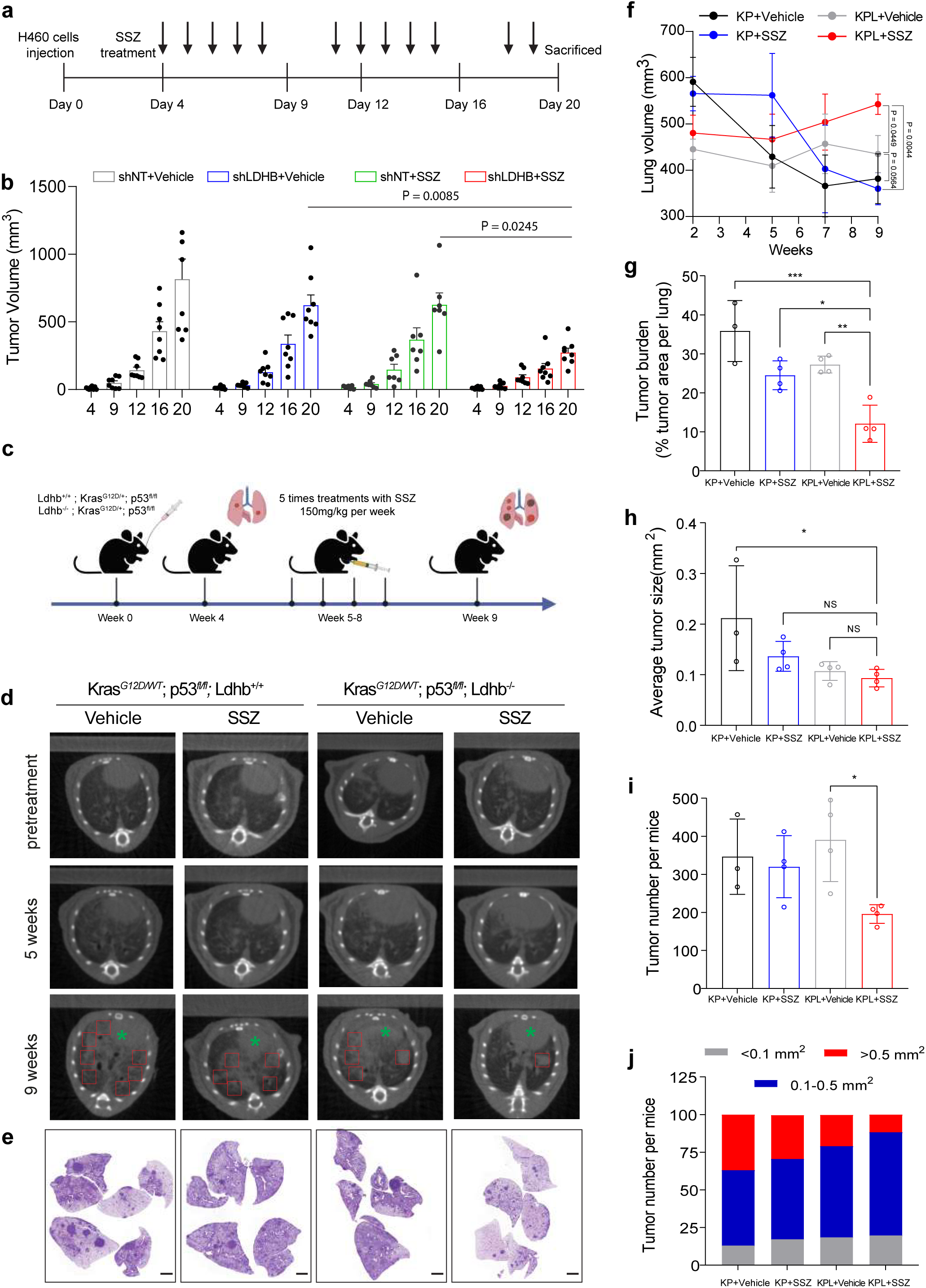
Combined inhibition of LDHB and SLC7A11 suppresses tumor growth in *Kras*-driven lung adenocarcinoma by inducing ferroptosis. **a,** The scheme of treatment in H460 xenograft tumor model with SSZ (150 mg/kg/day). **b,** Tumor volume of *KRAS*-mutant H460 xenografts stably expressing a control shRNA (shNT) or an LDHB-targeted shRNA (shLDHB) after treated with vehicle or SSZ (150 mg/kg/day). **c,** Schematic of autochthonous GEM models of *Kras*-induced adenocarcinoma with and without *Ldhb* deficiency. Established lung tumors in *LSL-Kras^G12D/WT^*; *p53*^fl/fl^ mice and *LSL- Kras^G12D/WT^*; *p53*^fl/fl^; *LDHB^fl/fl^* mice were treated with vehicle, SSZ (150 mg/kg/day) as indicated. **d,** Micro-CT images of tumors in *LSL-Kras^G12D/WT^*; *p53*^fl/fl^ mice and *LSL-Kras^G12D/WT^*; *p53*^fl/fl^; *LDHB^fl/fl^* mice at the indicated time points. Animals were scanned twice/week. **e,** H&E staining of lung tissue sections after 9-week treatment. Scale bar, 20 μM. **f-j,** Lung volume (f), lung tumor burden (g), average tumor size (h), tumor number (i) and tumor grade (number of nodules per mouse; j) after SSZ treatment. Data are shown as the mean ± s.d.. *p < 0.05. ** p < 0.01. ***p < 0.001 by two-way ANOVA. Ns, not significant.

Treatment in GEM models showed similar results: while SSZ exhibited only mild effect on tumor growth in KP mice (*Kras^G12D/WT^; p53^fl/fl^*), it markedly suppressed tumor development in KPL mice (*Kras^G12D/WT^; p53^fl/fl^; Ldhb^-/-^*), as evidenced by macroscopic examination of tumor lesions and histological evaluation of hematoxylin and eosin (H&E)-stained tissue sections (**Figure 5c, d, e**). Further scrutiny by quantitative measurement revealed that healthy lung volume was significantly preserved by SSZ treatment in KPL mice compared with KP mice (**Figure 5f**). Moreover, lung tumor burden (tumor area/total lung area), tumor size, and tumor number were significantly reduced by SSZ in KPL mice compared to KP mice (**Figure 5g, h, I, j**). These *in vivo* results, which corroborate *in vitro* cell death data, demonstrate that LDHB deficiency synergizes with SLC7A11 inhibition to suppress *KRAS*- dependent tumor growth by triggering ferroptosis.

### LDHB acts in parallel with NRF2 in regulating GSH synthesis and protecting against ferroptosis

The response of *KRAS*-dependent cell lines to ferroptosis induced by LDHB/SLC7A11 inhibition was heterogeneous, ranging from severe ferroptotic death to immediate onset of cell death and to complete resistance to ferroptosis (**Figure 2**), suggesting additional regulatory mechanisms acting in parallel with LDHB-mediated ferroptosis protection. NRF2 is the master antioxidant transcription factor that promotes ROS detoxification and *KRAS*-driven tumorigenesis by regulating the expression of SLC7A11, glutamate– cysteine ligase subunits GCLC and GCLM, and other effectors [11, 55]. We therefore examined whether the heterogeneous onset of ferroptosis is associated with NRF2 activity. Indeed, the expression of SLC7A11, but not of KEAP1, was associated with high expression of NRF2 in the NSCLC panel (**Supplementary Figure 10a**) [56]. Importantly, KRAS-dependent NSCLC cells that were hypersensitive to LDHB/SLC7A11 inhibition (A549, H460, H2122, H838, H2009, KP) exhibited high levels of NRF2 or NRF2 score based on prior studies [57, 58], whereas the outliers (H358, H23, and Calu-1), in which LDHB KD failed to sensitize the cells to Erastin or RSL3, are characterized by low NRF2 levels. The outliers also included *EGFR*-mutant (PC9, H1650) and *FGFR1-*amplified (H520) NSCLC and others (H1299, HT-1080), which could be explained by the fact that SLC7A11 was not affected by LDHB KD (**Supplementary Figure 10b, c, d**). These results suggest that NRF2 serves as a biomarker for predicting ferroptosis by targeting LDHB and SLC7A11 in *KRAS*-dependent cancer cells, consistent with previous finding that LDHB plays a unique role in KRAS-mutant lung cancer [27, 56].

To directly interrogate the role of NRF2 in LDHB-mediated ferroptosis protection, we forced the expression of NRF2 in H358 cells by genetic knockout (KO) of KEAP1 (**Figure 6a, b**). Compared to KEAP1 wildtype cells, KEAP1 KO H358 cells were highly sensitive to LDHB KD/Erastin- and LDHB KD/RSL3-induced ferroptotic death (**Figure 6c, d, e, f; Supplementary Figure 10e, f**), indicating that upregulation of NRF2 is required and sufficient for the hypersensitivity of *KRAS*-mutant cancer cells to ferroptosis induced by the combined inhibition of LDHB and SLC7A11 or GPX4.

**Figure 6.**
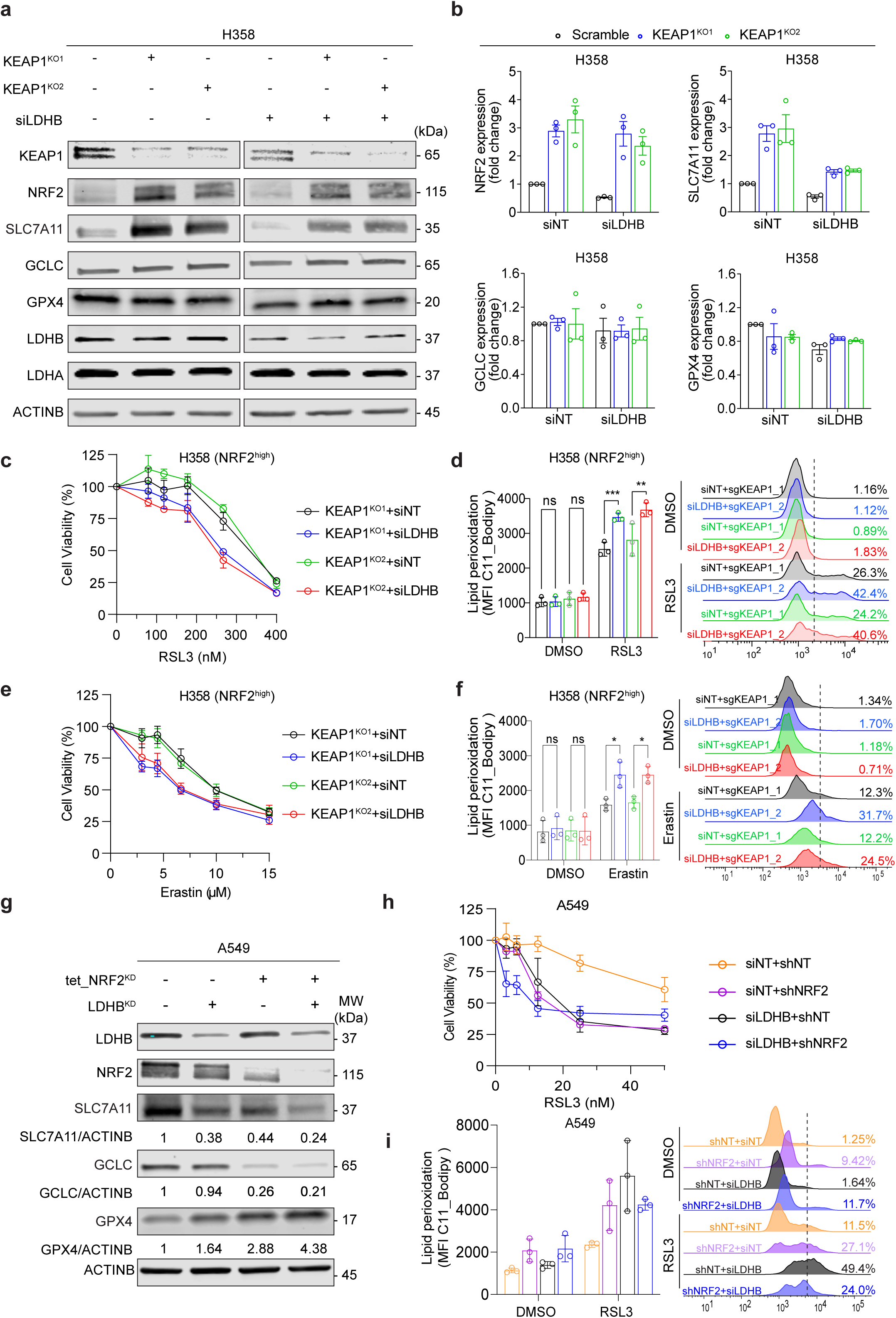
LDHB acts in parallel with NRF2 to regulate the SLC7A11/GSH/GPX4 axis. **a,** Immunoblots of H358 cells transduced by and stably expressing sgControl (pXPR_BRD003_Luci) or KEAP1-targeted sgRNAs (sgKEAP1_1, sgKEAP1_2) after transfected with siNT or siLDHB. **b,** Fold changes of NRF2, SLC7A11, GCLC, and GPX4protein levels in the indicated cells. Quantification was based the results in **a**. **c,** Viability assay of H358 cells transduced as in **a** and treated with RSL3 for 24 h. **d,** Lipid peroxidation of H358 cells transduced as in **a** and treated with RSL3 for 6 h. **e,** Viability assay of H358 cells transduced as in **a** and treated with Erastin for 24 h. **f,** Lipid peroxidation of H358 cells transduced as in **a** and treated with Erastin for 10 h. **g,** Immunoblots of A549 cells stably expressing a doxycycline (0.5 μM)-inducible control shRNA (shContorl) or a doxycycline (0.5 μM)-inducible shRNA against NRF2 (shNRF2) after transfected with either siNT or siLDHB for 72 h. **h,** Viability assay of A549 cells transduced and induced as in **g** and treated with RSL3 for 72 h. **i,** Lipid peroxidation of A549 cells transduced and induced as in **g** and treated with RSL3 for 6 h.

**Figure 7.**
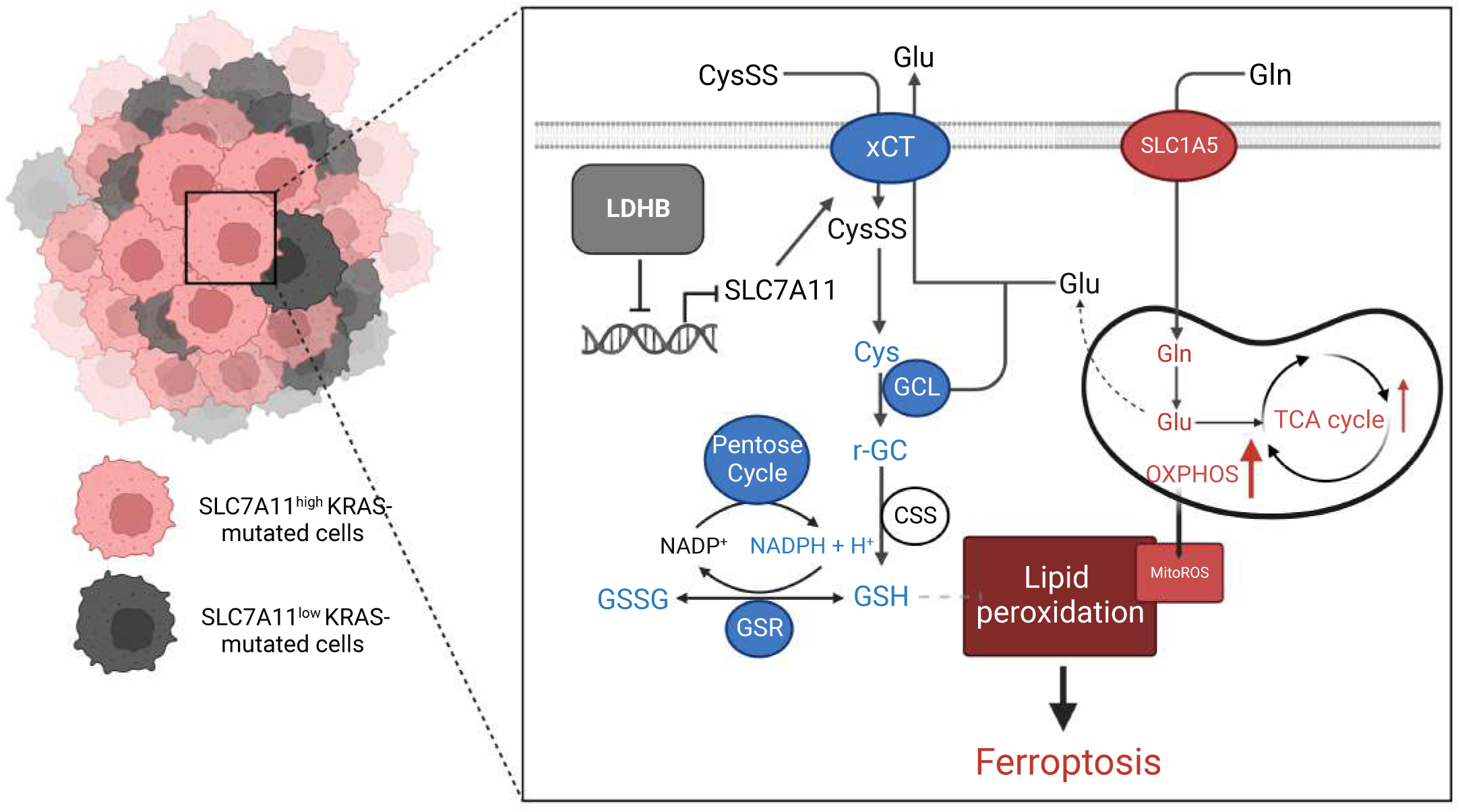
Working model illustrating a non-canonical role of LDHB in ferroptosis surveillance in *KRAS*-driven lung cancer. LDHB plays a critical but non-canonical role in protecting *KRAS*-driven lung cancer against ferrotoptosis. Specifically, LDHB upregulates the expression of SLC7A11 in an NRF2-independent manner, thereby promoting GSH synthesis. Suppression of LDHB evokes oxidative stress and synergizes with ferroptosis inducers that block SLC7A11 and GPX4, key antioxidant hubs whose activity is exquisitely dependent on GSH production and utilization. As a result, combined inhibition of LDHB and SLC7A11 or GPX4 synergistically disrupts GSH-mediated interception of ferroptosis, which promotes glutamine entry into the mitochondria, increases OXPHOS and mitoROS production, and ultimately translates into glutaminolysis-dependent ferroptosis.

In light of the fact that LDHB and NRF2 converge in regulating SLC7A11 expression, we investigated whether their inhibition synergizes in ferroptosis induction. Inducible suppression of NRF2 (tet_NRF2^KD^) decreased the expression of SLC7A11 but not of LDHB, as did LDHB KD, which reduced SLC7A11 but barely affected NRF2 expression (**Figure 6g**). Simultaneous KD of NRF2 and LDHB further reduced SLC7A11 and the downstream GCLC and primed sensitivity to the ferroptosis inducer RSL3 (**Figure 6g, h**). These results suggest that LDHB acts in parallel to NRF2 in ferroptosis protection in *KRAS*-dependent lung cancer cells.

## DISCUSSION

In this study, we uncover that the glycolytic enzyme LDHB mediates an unexpected mechanism of ferroptosis surveillance in *KRAS*-dependent lung cancer. Metabolic, genetic and biochemical evidence accommodates a model in which LDHB is a critical regulator of the SLC7A11/GSH/GPX4 antioxidant program and LDHB deficiency promotes the hypersensitivity of *KRAS*-dependent cancer cells to ferroptosis inducers. As a result, the combination of LDHB inhibition with ferroptosis inducers synergistically suppresses *KRAS*-dependent cancer cells and tumors by inducing lethal ferroptosis *in vitro* and *in vivo*. We further demonstrate that metabolic synthetic lethality of SLC7A11 inhibition in LDHB-deficient cancer cells is fueled by increased glutaminolysis and subsequent pro-ferroptotic metabolic activities in mitochondria. These findings reveal a novel oncogene-specific mechanism of ferroptosis defense and suggest a previously unrecognized strategy to selectively target *KRAS*-mutant lung cancer.

Oncogenic KRAS deregulates a variety of cellular processes to promote tumorigenesis. This includes profound alterations of metabolic pathways to meet the increased energy, biosynthesis, and redox requirements of cancer cells [59, 60]. In particular, abnormal ROS production, a functional requirement for KRAS-mediated tumorigenicity [10], is one of the major metabolic manifestations in *KRAS*-driven cancer. To achieve redox balance, cancer cells have evolved complex antioxidant programs that leverage ROS levels to favor tumor growth but prevent cell death [12, 15, 37]. However, the mechanisms underlying the antagonism between cancer metabolism that promotes oxidative stress and the ROS surveillance mechanisms that are exploited by mutant KRAS, which may open therapeutic opportunities for cancer treatment, remain inadequately understood [34]. Here, we report for the first time an antioxidant function for LDHB in protecting *KRAS*-mutant lung cancer from ferroptosis, a ROS-dependent non-apoptotic cell death program triggered by iron-dependent accumulation of lipid peroxides in plasma membranes and intracellular organelles such as mitochondria [31]. This finding is consistent with the increasingly appreciated consensus that bypass or inactivating ferroptosis is a hallmark mechanism co-opted by oncogenic KRAS mutations to overcome the oxidative barrier during tumor development, progression, and metastasis [40-42, 61, 62].

LDHB has a unique role in *KRAS*-driven lung adenocarcinomas [27, 28], but the underlying mechanism remains poorly understood. Here, we find that LDHB promotes GSH synthesis and defends against ferroptosis by upregulating SLC7A11, the functional subunit of the cystine/glutamate antiporter system Xc^−^ (SLC7A11 and SLC3A2), which takes up extracellular cystine to synthesize the antiferroptotic metabolite GSH [36, 63], a cofactor of GPX4 that neutralizes lipid peroxides and suppresses ferroptosis [33]. SLC7A11 and GPX4 form the main engine for GSH biosynthesis and utilization, and the SLC7A11/GSH/GPX4 axis is the major surveillance system for defense against ROS-dependent ferroptosis in cancer cells [31, 35, 64, 65]. Previous studies have indicated a functional link between SLC7A11 and mutant KRAS: SLC7A11 is overexpressed in *KRAS*-mutant cancers and has prognostic significance; inactivation of SLC7A11 leads to selective damage in cancer cells harboring mutant KRAS [39] and SLC7A11 promotes *KRAS*-induced tumorigenicity by increasing glutathione synthesis [63, 66]. However, the metabolic processes and context-specific factors that modulate ferroptosis sensitivity of *KRAS*-dependent lung cancer by impinging on SLC7A11 are not fully known. Our results extend the prior work and suggest that SLC7A11 is a key effector of LDHB and contributes to *KRAS*-dependent lung cancer through its cooperative function in ROS scavenging and ferroptosis surveillance. Our finding that LDHB is a key component of ROS surveillance mechanisms in *KRAS*-driven cancer is also consistent with the notion that LDHB plays an important role in homeostasis regulation including autophagy [26, 67].

LDHA and LDHB are subunits of the tetrameric enzyme LDH. While LDHA catalyzes the reduction of pyruvate to lactate, LDHB oxidizes lactate to pyruvate, coupled with NAD^+^ to NADH [23]. Despite different distributions and functions, both LDHA and LDHB are important for ATP generation and homeostasis under physiologically low O2 (anaerobic glycolysis) and also for the reprogramed cancer metabolic state of aerobic glycolysis known as the Warburg effect [24, 25]. LDHA and LDHB are predominantly distributed in cytoplasm and mitochondria, respectively, but they can also be found in the nucleus where they binds to mRNA, suggesting that LDHB has additional roles in post-transcriptional modification of gene expression independent of its canonical function in glucose metabolism [68]. Indeed, LDHA has been shown to promote T cell effector functions by increasing acetylation and transcription of INF-γ [69]. Our finding that LDHB silencing activates STAT1, a key effector of the INF-γ/JAK pathway, which suppresses the expression of SLC7A11 is plausibly consistent with the notion that LDH has additional roles beyond glycolysis [23], although the possibility that lactate rather than LDHB itself modulates INF-γ/JAK pathway activity cannot be completely excluded [67]. Whether the enhanced expression of LDHB and the newly identified role of LDHB in regulating SLC7A11-mediated GSH synthesis in *KRAS*-dependent lung cancer, which favors tumor growth, is due to its enzymatic or its gene regulatory function requires further investigation.

We further show that increased glutaminolysis underpins the metabolic synthetic lethality of LDHB deficiency and ferroptosis inducers, as evidenced by the observation that challenges to the SLC7A11/GSH/GPX4 axis in LDHB-deficient cancer cells lead to acute activation of glutamine uptake and glutaminolysis. Glutaminolysis is a mitochondrial process in which glutaminases (GLS) catalyze the conversion of glutamine into glutamate [70], and is an important pathway that intersects with glycolysis, the TCA cycle, redox homeostasis, and lipid and amino acid homeostasis. The overwhelming metabolic stress and ensuing ferroptosis incurred by LDHB depletion and ferroptosis inducers as a consequence of glutaminolysis is consistent with previous observations. Glutaminolysis positively regulates aerobic energy production in mitochondria, increases lipid ROS production and promotes ferroptosis [54]. Importantly, several studies have showed that inhibiting SLC7A11 activities suppresses glutamine-derived glutamate export, enhance glutamate to replenish the TCA cycle intermediates, and promote ROS production [63, 71, 72], suggesting that SLC7A11 antagonizes pro-ferroptotic glutaminolysis and that increasing glutamine metabolism under blockage of GSH synthesis upregulates ROS levels and promotes ferroptosis, which may provide a novel treatment guideline option for ferroptosis-based tumor therapy.

In light of the concept that mutant KRAS alters metabolic pathways and that contextually activated metabolic alterations confer selective vulnerabilities, exploiting metabolic synthetic lethality is increasingly appreciated as a strategy to target *KRAS*-mutant lung cancer [39-41, 73, 74]. As metabolic ROS is causally linked to mutant KRAS-induced tumorigenicity and requires homeostatic mechanisms to maintain ROS levels within a threshold favorable for tumor development (14,27-30,41), the identification of LDHB and SLC7A11/GSH/GPX4 converging on a role as ROS scavengers reveals an unanticipated metabolic vulnerability in *KRAS*-mutant cancers. We further show that KEAP1-mutant or NRF2-high cancer cells are dependent on LDHB and increased SLC7A11/GSH/GPX4 activity, and this property can be therapeutically exploited through the pharmacological inhibition using ferroptosis inducers [9]. NRF2 is the master transcriptional regulator of endogenous antioxidant synthesis that has been shown to be induced by oncogenic KRAS and its effector pathways [11]. Our finding that LDHB and NRF2 independently regulate the expression of SLC7A11 is in line with previous studies reporting cancer subtype-specific alterations in glycolysis and GSH biogenesis. Future studies will be necessary to clarify whether and how LDHB contributes to ferroptosis surveillance in *KRAS*-mutant cancer harboring wildtype KEAP1 and expressing low levels of NRF2.

In conclusion, we discover that LDHB mediates ferroptosis resistance in *KRAS*-dependent lung cancer by promoting SLC7A11-mediated GSH metabolism and that the mechanistic underpinning of the ferroptosis phenotype induced by SLC7A11 inhibitors in LDHB- deficient cells involves activation of pro-ferroptotoc glutaminolysis. Our results provide strong mechanistic support for the combining the inhibition of LDHB with ferroptosis inducers by targeting the SLC7A11/GSH/GPX4 antioxidant nexus for treating KRAS-dependent lung cancer and pinpoint a biomarker that could be used to identify patients likely to benefit from this combination treatment. In addition, our study supports the rationale for the development of LDHB-specific inhibitors for the clinical implementation of our findings and provides further evidence that targeting oncogene-specific ferroptosis mechanisms may facilitate the identification of rational strategies for the treatment of KRAS-driven lung cancer.

## METHODS

### Cell culture and reagents

*KRAS*-mutant cancer cell lines used in this study (**Supplementary Table S3**) were obtained from American Type Culture Collection (ATCC, Manassas, VA, USA). BEAS-2B and murine KP cells derived from a GEM model (*Kras^G12D^*;*Trp53*^−/−^) were described previously [75]. Cells were cultured in RPMI-1640 medium or Medium 199 (Cat. #8758 and #4540; Sigma-Aldrich, St. Louis, MO, USA) supplemented with 10% fetal bovine serum/FBS (Cat. #10270-106; Life Technologies, Grand Island, NY, USA) and 1% penicillin/streptomycin solution (Cat. #P0781, Sigma-Aldrich). The cells were authenticated by DNA fingerprinting and confirmed free from mycoplasma contamination (Microsynth, Bern, Switzerland). All inhibitors used in this study were listed in **Supplementary Table 4**.

### Cell viability, cell death and clonogenic survival assay

Cells seeded in 96-well plates (2500 cells/well) were dosed 24 h later with different inhibitors for 72 h or indicated time. Cell viability was determined by APH assay as previously described (14, 31). The efficacy of drugs on cell growth was normalized to untreated control. Each data point was generated in triplicate and each experiment was done three times (n=3). Best-fit curve was generated in GraphPad Prism [(log (inhibitor) vs response (-variable slope four parameters)]. Error bars are mean ± SD.

To measure cell death, cells were seeded in 6-well plates for overnight culture. Cell death was determined using SYTOX dead cell stain sampler Kit (ThermoFisher Scientific, S34862) according to the manufacturer’s instructions. Cell number counting was determined by Countess™ 3 (ThermoFisher Scientific) according to the manufacturer’s instructions.

Clonogenic assay was done as we described previously [76]. In brief, cells seeded in 6- well plates (3000 cells/well) were dosed 24h later and continually treated with rapamycin for 7 days (refresh drugs every three days), the resulting colonies were stained with crystal violet (0.5% dissolved in 25% methanol).

### Gene silencing by small interfering RNA (siRNA), short hairpin RNAs (shRNA) and single-guide RNAs (sgRNA)

Transient knockdowns were mediated by siRNAs. Cells cultured in triplicate at 50-70% confluency were transfected with specific pooled siRNA Oligo Duplex (Origene Technologies, Rockville, MD, USA) or control siRNA Duplex. Transfection with Lipofectamine 2000 transfection reagent (Invitrogen, 11668027) was performed according to the manufacturer’s protocol. Stable knockdown was achieved via lentiviral constructs expressing a specific shRNA or a scramble shRNA (negative control). Recombinant lentiviruses were produced by transfecting HEK293T cells with pCMV-VSV-G (VSV-G protein), pCMV-dR8.2 (lentivirus packaging vector) and shRNA-expressing constructs using Lipofectamine 2000 transfection reagent as previously described. The supernatant containing lentiviruses was collected, filtered through 0.45 µM filters, and stored in aliquots at -80°C, or immediately used to infect recipient cells. After infection, cells were selected in puromycin (1.5 µg/ml) and further passaged in culture for functional assays. A full list of shRNA and siRNA sequences is provided in **Supplementary Table 3**.

### RNA isolation, quantitative real-time PCR (qRT-PCR) and RNA sequencing

Total RNA was isolated and purified using RNeasy Mini Kit (Qiagen, Hilden, Germany). Complementary DNA was synthesized by the High capacity cDNA reverse transcription kit (Applied Biosystems, Foster City, CA, USA) per manufacturer’s instructions. Real-time PCR was performed in triplicate on a 7500 Fast RealTime PCR System (Applied Biosystems) using TaqMan primer/probes (Applied Biosystems) shown in **Supplementary Table 3.** Normalization was based on the ΔΔCT method.

Total RNA was isolated and purified with RNeasy Mini Kit (Qiagen, 74106). The quality control assessments, generation of libraries, and sequencing were conducted by the Next Generation Sequencing Platform, University of Bern as previous described. The quality of the RNA-seq data was assessed using fastqc v. 0.11.9 (Andrews 2022) and RSeQC v. 4.0.0 (Wang, Wang, and Li 2012). The reads were mapped to the reference genome using HiSat2 v. 2.2.1 (Kim, Langmead, and Salzberg 2015). FeatureCounts v. 2.0.1 (Liao, Smyth, and Shi 2014) was used to count the number of reads overlapping with each gene as specified in the genome annotation (Homo_sapiens. GRCh38.104). The Bioconductor package DESeq2 v1.36.0 (Love, Huber, and Anders 2014) was used to test for differential gene expression between the experimental groups. ClusterProfiler v4.4.4 (Wu et al. 2021) was used to identify gene ontology terms containing unusually many differentially expressed genes. Gene set enrichment analysis (GSEA) (Subramanian et al. 2005) was run in ClusterProfiler v4.4.4 (Wu et al. 2021) using gene sets from KEGG (Kanehisa, Sato, and Kawashima 2022) and MSigDb (Liberzon et al. 2015). An interactive Shiny application v1.6.0 (Chang et al. 2022) was set up to facilitate the exploration and visualisation of the RNA-seq results. All analyses were run in R (version 4.2.0; R Core Team 2022).

### Immunoblotting, immunohistochemistry and immunofluorescence

Cell lysates were prepared and western blot analysis was performed as described (14,31). In brief, equal amounts of protein lysates resolved by SDS-PAGE (Cat. #4561033; Bio-Rad Laboratories, Hercules, CA, USA) and transferred onto nitrocellulose membranes (Cat. #170-4158; Bio-Rad). Membranes were then blocked with blocking buffer (Cat. #927-4000; Li-COR Biosciences, Bad Homburg, Germany) for 1 h at room temperature (RT) and incubated with appropriate primary antibodies overnight at 4 °C (**Supplementary Table 3**). IRDye 680LT-conjugated goat anti-mouse IgG (Cat. #926- 68020) and IRDye 800CW-conjugated goat anti-rabbit IgG (Cat. #926-32211) from Li-COR Biosciences were used at 1:5000 dilutions. Finally, signals of membrane-bound secondary antibodies were imaged using the Odyssey Infrared Imaging System (Li-COR Biosciences).

For immunofluorescence, tumor cells grown on poly-lysine-treated coverslides were fixed with 4% paraformaldehyde for 15 min at RT and permeabilized with cold methanol (−20 °C) for 5 min or with 0.1% Triton X-100/PBS at RT for 15 min before incubated overnight at 4 °C with primary antibodies (**Supplementary Table 3**). The cells were incubated for 1 h at RT with Alexa Fluor 647 goat anti-mouse IgG (Cat. #<u>A21236</u>) or Alexa Fluor 488 goat anti-Rabbit IgG (Cat. #<u>A11034</u>) from Invitrogen (Eugene, OR, USA). Nuclei were counterstained by 4′,6-diamidino-2-phenylindole. Images were acquired on a ZEISS Axioplan 2 imaging microscope (Carl Zeiss MicroImaging, Göttingen, Germany) and processed using Adobe illustrator CC 2017 (Adobe Systems, San Jose, CA, USA).

Immunohistochemical study was performed as we described previously (31,32). In brief, surgically removed xenograft tumors were formalin-fixed and paraffin-embedded (FFPE). FFPE tumors were sectioned at 4 μm, deparaffinized, rehydrated and subsequently stained with hematoxylin and eosin (H&E) and appropriate antibodies (**Supplementary Table 3**) using the automated system BOND RX (Leica Biosystems, Newcastle, UK). Visualization was by the Bond Polymer Refine Detection kit (Leica Biosystems) as instructed by the manufacturer. Images were acquired by PANNORAMIC® whole slide scanners, processed by Case Viewer (3DHISTECH Ltd.). The staining intensities of the whole slide (two tumors/group) were quantified by QuPath software.

### Immunofluorescence microscopy

The cells were treated with indicated conditions and then stained with C11-Bodipy for 30 mins in cell culture medium. The cells were washed with PBS for 5 min three times. ProLong Diamond anti-fade mountant with DAPI (ThermoFisher, P36962) was added to stain the nucleus. All images were captured on a fluorescence microscope at the indicated objective with constant laser intensity for all samples.

### LS-MS, labelling of D-glucose-1,2-[13C2] and metabolomics data analysis

Cells seeded in six-well plates were cultured overnight with 2 mL fresh medium with or without 25 mM D-glucose-1,2-[13C2] (Sigma-Aldrich, 453188) before treatment. Cells were then washed twice with PBS and solvent (75 mM ammonium carbonate, pH was adjusted to 7.4 with acetic acid). Pre-cooled (− 20 °C) extraction solvent (40% acetonitrile, 40% methanol, 20% nanopure water) was immediately added to the plates. Cells were then scraped from the dish on the ice, vortexed for 30 second and immediately stocked in – 20 ℃ for 1 hour and then transferred to – 80 ℃. LC–MS measurement and analysis were described previously. Metaboliomic data analysis with MetaboAnalyst 5.0 software.

### OCR measurements

Cells were seeded overnight in Seahorse XF96 V3PS cell culture microplates (Agilent Technologies, 101085-004) and allowed cell attached the plates and then treated with indicated drugs for indicated time. The XF sensor cartridges were hydrated overnight with sterile ddH2O in a CO2-free incubator at 37 °C. Cells were washed twice and changed into Seahorse XF DMEM or RPMI medium (Agilent Technologies, 103680-100, 103681-100) containing 10 mM glucose, 0.5 mM pyruvate, 2 mM glutamine adjusted to pH 7.4. Then the cells were incubated in a CO2-free incubator for 1 h. 1 μM oligomycin, 1.0 μM and 1.5 μM FCCP, a mixture of 1 μM rotenone and 1 μM antimycin A were added successively. A full list of chemicals is listed in Supplementary Table. The data were analyzed with Seahorse Wave (Agilent Technologies) and normalized to 50 ng DNA, which is quantified by CyQUANT™ Cell Proliferation Assay kit (Thermo Fisher Scientific, C7026) according to the manufacturer’s protocol.

### Lipid peroxidation and MitoSOX measurements

Cells were seeded in 6-well plates or 12-well plates at 0.1 to 0.3 million or 0.05 to 0.2 million cells per well overnight before treatment, respectively. For lipid peroxidation measurements, Cells were stained 2.5 μM C11_BODIPY (D3861, Invitrogen) for 30 min at 37 °C in cell culture medium after treatment. Then cells were washed with PBS, detached with trypsin and centrifuged two times (3 min, 400 g). ROS measurement was performed as described previously. MitoSOX measurement using MitoSOX™ Red Mitochondrial Superoxide Indicator (M36008, ThermoFisher Scientific). Cells were washed with PBS, detached with trypsin and centrifuged (5 min, 400 g), resuspended and covered with 5 μM working solution of MitoSOX reagent in HBSS and incubated at 37 °C for 10 min in FACS tubes. Wash cells gently with warm HBSS buffer three times and then prepare cells for detecting and analyzing. Flow cytometry determined all the samples following a standard procedure with a FACS lyric instrument (BD Biosciences). Data were analyzed using the FlowJo V10 workspace software.

### *In vivo* mouse study

Mouse studies were conducted in accordance with Institutional Animal Care and Ethical Committee-approved animal guidelines and protocols. Implantable tumor models were performed in age- and gender-matched NSG (NOD-*scid IL2Rγ^null^*) as we previously described (31,32). Specifically, 1 × 10^6^ A549, H838 and H460 cells stably expressing shLDHB or a control shRNA were 1:1 mixed with BD Matrigel Basement Membrane Matrix (Cat. #356231; Corning, NY, USA) and subcutaneously inoculated in left and right flanks (0.5x10^6^/injection). When tumors were palpable, mice were randomly assigned to treatment groups. Tumors were measured every three days, with their size calculated as follows: (length × width^2^)/2. For survival analysis, the mice were closely monitored on a daily basis, and the size of tumors was measured with a caliper every 4-5 days. Mice were sacrificed when the tumor volume reached 1000 mm^3^.

*Kras^LSL-G12D/WT^;p53^flox/flox^*; *Ldhb*^+/+^ (KP) mice and KrasLSL-G12D/WT;p53flox/flox; Ldhb−/− (KPL) mice have been described [20, 28]. Two weeks after infection with AAV-Cre virus. all mice were scanned with microCT (X-RAD SmART–Precision X-Ray) to determine the basal line of lung tumor volumes. Lung volume by 3D Slicer software. Mice were scanned with microCT every two times a week to assess tumor development and sacrificed at the week end of experiments. The microCT images were processed and analyzed with Fiji and 3D Slicer version 4.13 according to a previously published protocol.

### Statistical analysis

Statistical analyses were performed using GraphPad Prism 7.01 (GraphPad Software Inc., San Diego, CA, USA) unless otherwise indicated. In all studies, data represent biological replicates (n) and are depicted as mean values ± SD or mean values ± SEM as indicated in the figure legends. In all analyses, *P* values less than 0.05 were considered statistically significant. For the survival analysis, patients were grouped by gene expression, where ‘high’ and ‘low’ expression groups were stratified by the optimal cut-off value.

## Data availability

The RNA sequencing raw data have been deposited at GEO under the accession code: GSE224098. The metabolomics data and other data supporting this study are available within the Source Data File. All other data is available in the Article and Supplementary Information. Source data are provided with this paper.

## Acknowledgements

We acknowledge Christelle Dubey (Laboratory of General Thoracic Surgery, Inselspital, Bern University Hospital, University of Bern) for animal studies and technical support, and Translational Research Unit at Institute of Tissue Medicine and Pathology, University of Bern for assistance of IHC. We thank the science technology platforms and core facilities at Department of BioMedical Research (DBMR) and University of Bern including the Next Generation Sequencing (NGS) Platform, the Flow Cytrometry and Cell Sorting Facility, the Live Cell Imaging Core Facility, the Microscopy Imaging Center, the Experimental Animal Center EAC of the University of Bern. This work was supported by grants from Swiss National Science Foundation (SNSF #310030_192648 to R.-W.P.) and the Swiss Cancer Research Foundation (#KFS-4851-08-2019 to R.-W.P.), and PhD fellowship from China Scholarship Council (to L.Z., J.Z. and H.Z.).

## Authors’ contributions

L.Z. conceived the project, designed and performed the experiments, analyzed the data and wrote the paper. H.D. generated the stable LDHB knockdown cell lines and the Ldhb- deficient mouse model. J.Z. helped with FACS analysis and in vivo mouse studies. N.Z. performed the metabolomics studies and data analysis. G.A.G. and R.B. contributed to RNA sequencing data analysis and statistical analysis. H.Y., Z.Y., G.Y., D.X., and H.Z. assisted with in vitro and in vivo studies and helpful discussions. Q.Z. and R.A.S. provided resources. T.M.M. contributed to the study with Ldhb-deficient mouse model and acquired grant support. P.D. provided resources and financial support. RWP conceptualized the project, supervised the study, acquired grant support and wrote the paper. All authors contributed to manuscript revision and review.

