## Supplementary Figures for "A non-canonical function of LDHB promotes SLC7A11-mediated glutathione metabolism and protects against glutaminolysis-dependent ferroptosis in *KRAS*-driven lung cancer"

Supplementary Figure 1

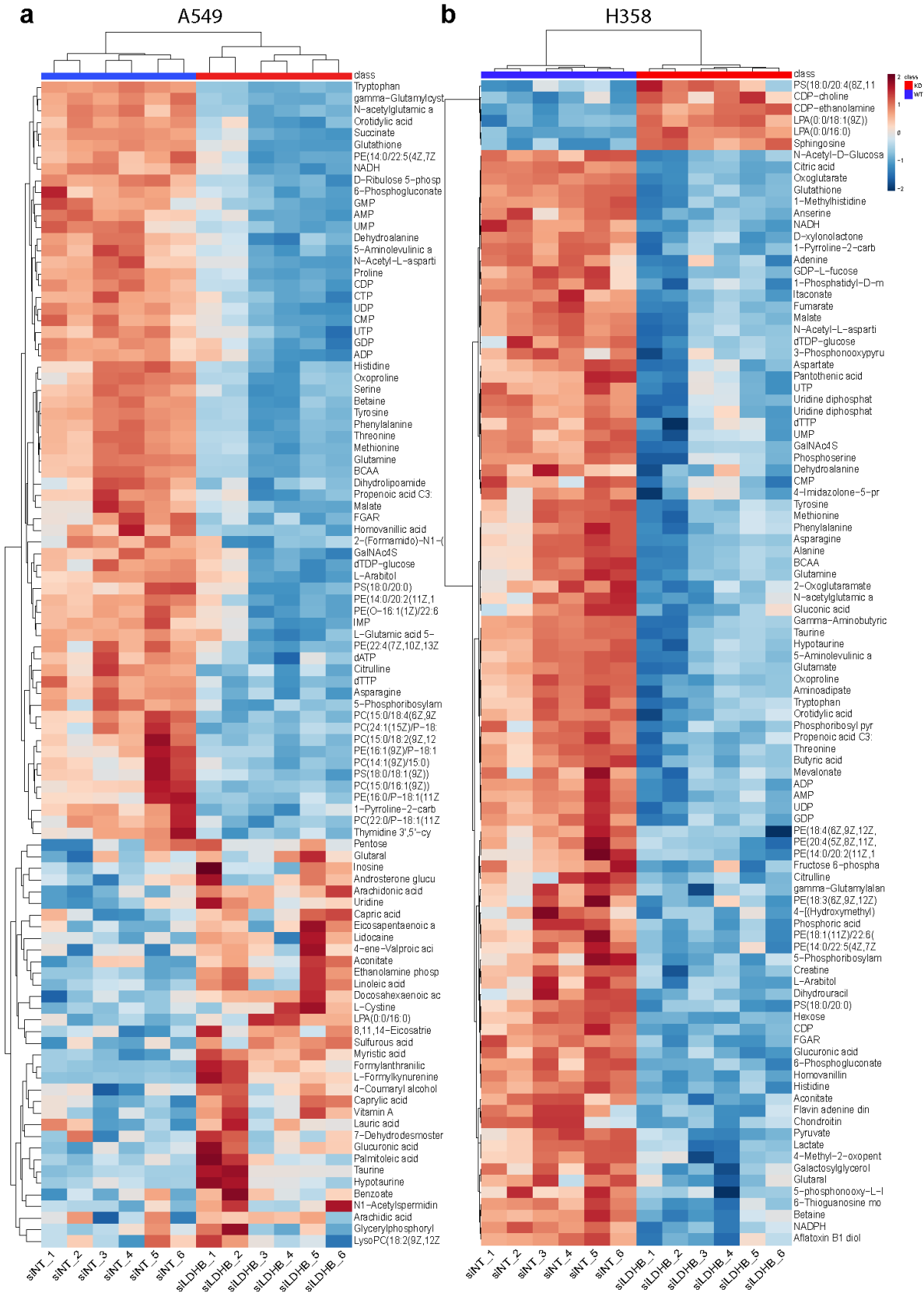

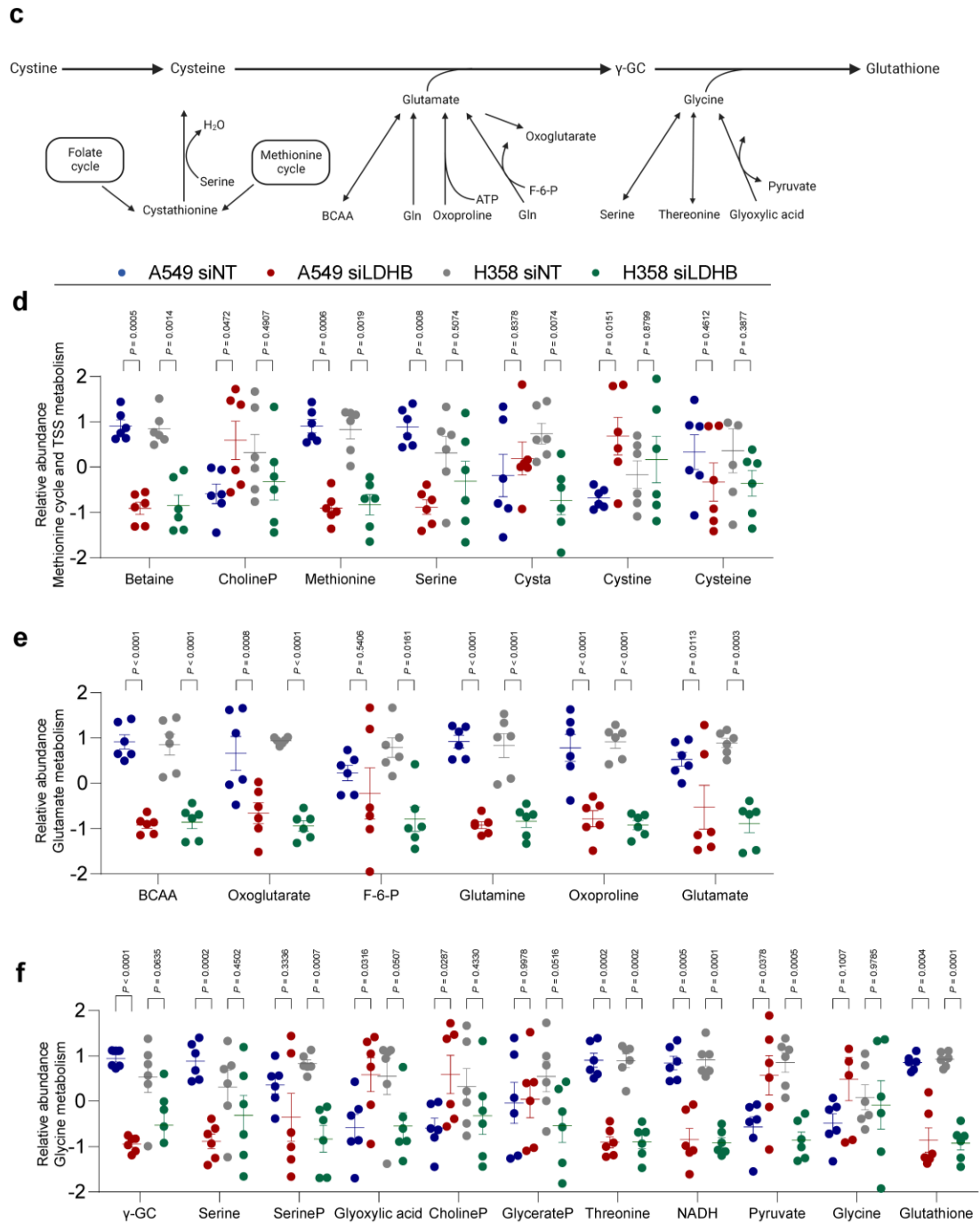

**Figure S1, Metabolomics analysis revealed that LDHB regulates GSH biosynthesis.** a-b, heat maps showing the top 100 significantly different metabolites in siLDHB A549 and H358 cells compared to siNT cells. Metabolomics profiling was analyzed 48 h post transfection. c, schematic illustrating methionine cycle, TSS pathway, cysteine biosynthetic pathway, glutamate metabolism, glycine metabolism contributing to GSH synthesis. d, e, f, normalized abundance of the indicated metabolites in the indicated samples, with the statistical analysis by two-way ANOVA.

Supplementary Figure 2

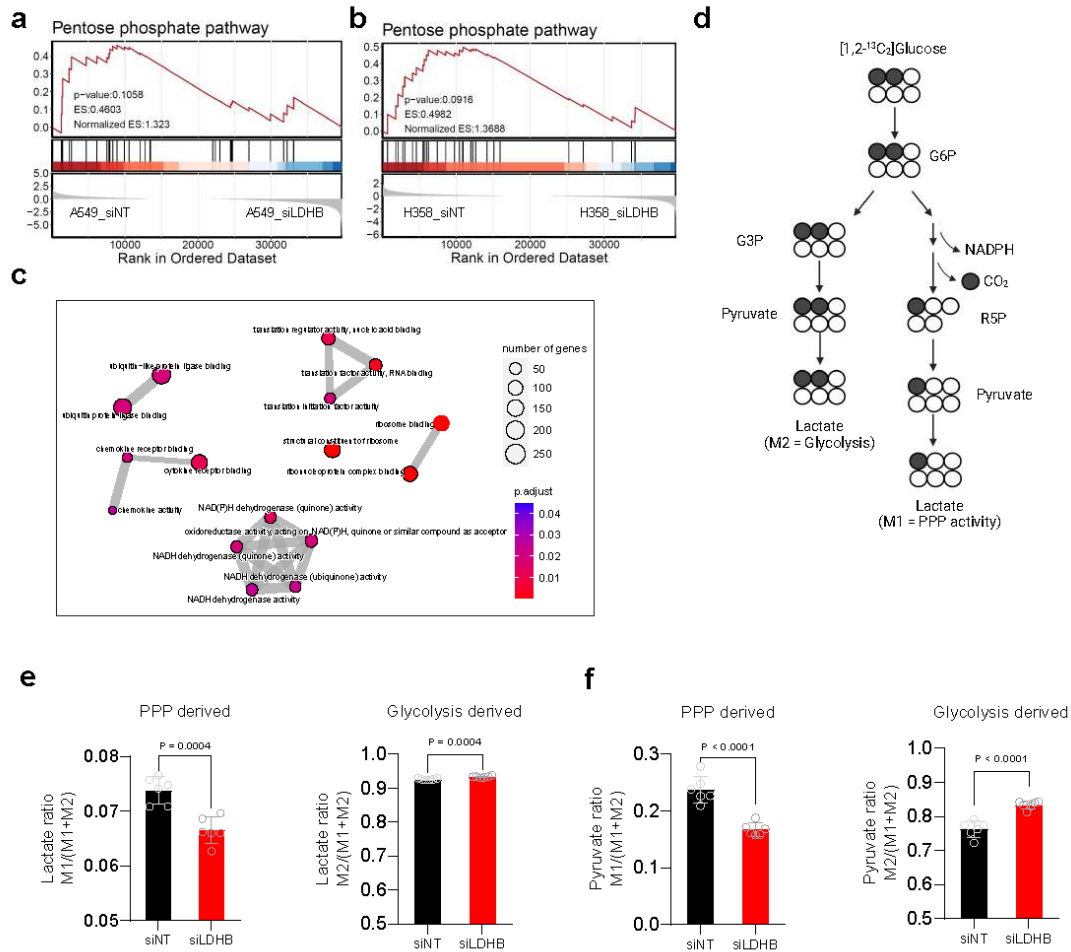

**Figure S2, LDHB silencing impairs the pentose phosphate pathway (PPP) and NADPH synthesis.** a-b, GSEA plots of the PPP signature in the indicated groups (siNT A549 vs. siLDHB A549; siNT H358 vs. siLDHB H358). c, an emap plot showing the NADPH bio-synthesis pathway enriched in siNT A549 compared to siLDHB A549 cells. d, schematic of [1,2-<sup>13</sup>C<sub>2</sub>] glucose metabolism through glycolysis and PPP pathway. e-f, A549 cells transfected with siNT or siLDHB for 48 hours were subjected to analysis of lactate and pyruvate derived from [1,2-<sup>13</sup>C<sub>2</sub>] glucose.

related to Fig 1

##### Supplementary Figure 3

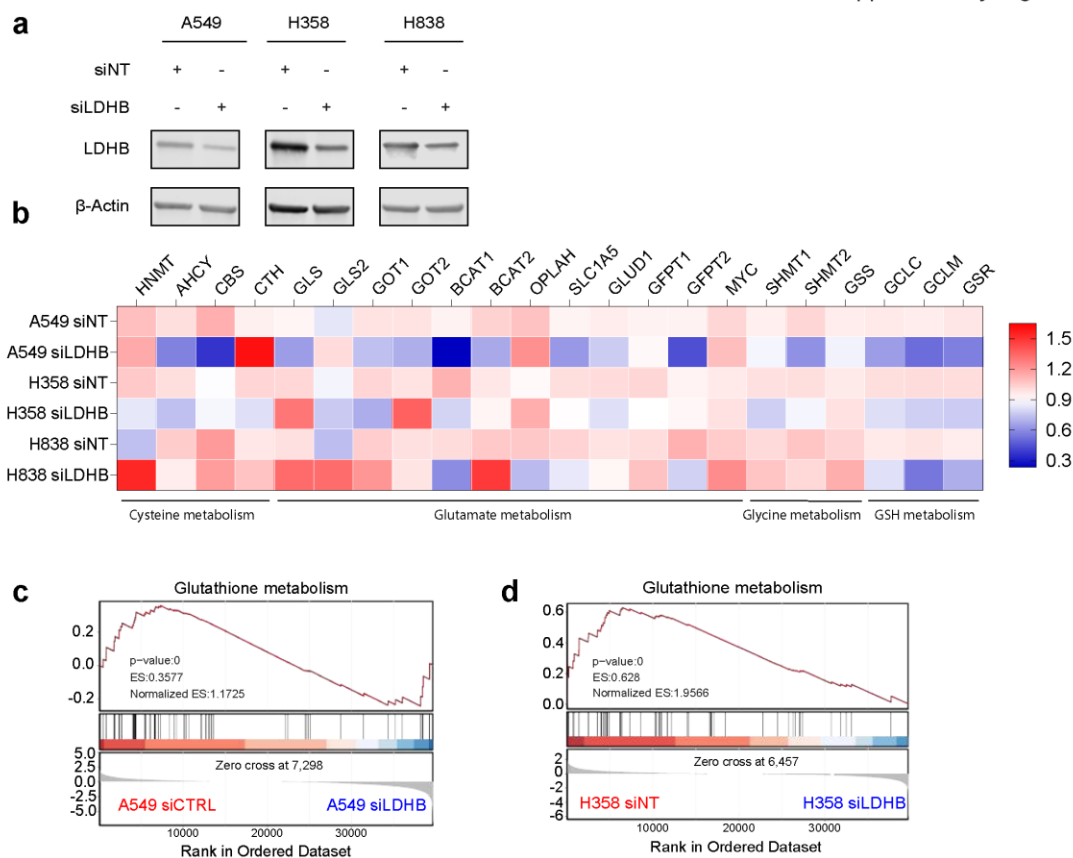

**Figure S3, RNA sequencing analysis revealed that LDHB regulates GSH biosynthesis.** a, Immunoblots of LDHB protein levels in the indicated cells. b, heat map showing the fold change of expression of the indicated genes involved in GSH synthesis (refer to Fig. 1c). The siLDHB group was normalized to the siNT group, with each square representing the mean value of three independent experiments (n=3). c-d, GSEA plots of GSH metabolism signatures A549 and H358 cells with and without LDHB KD.

related to Figure2

### Supplementary Figure 4

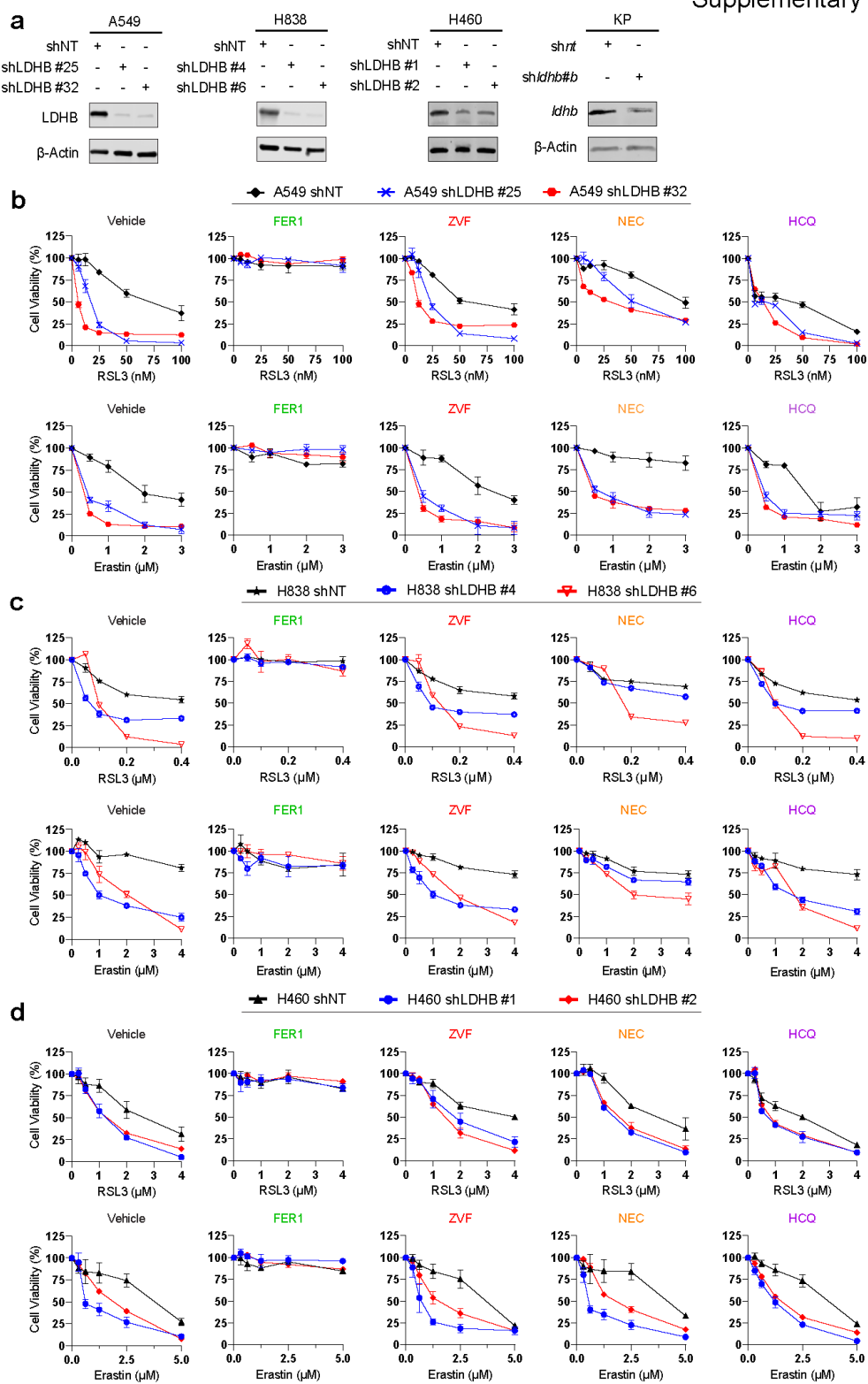

Supplementary Figure 4

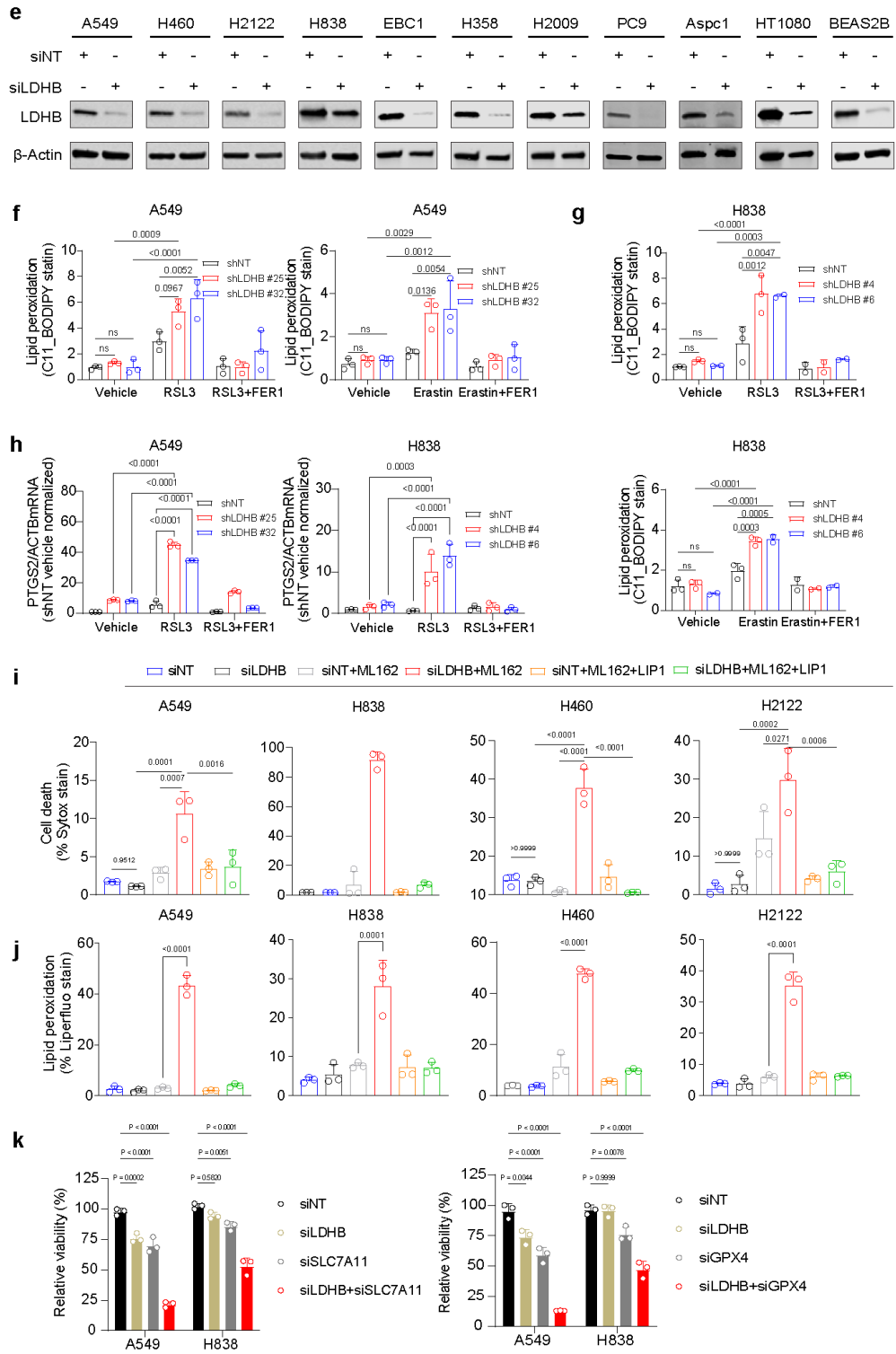

**Figure S4, LDHB inhibition sensitizes KRAS-dependent lung cancer cells to ferroptosis inducers.**

a, immunoblot analysis of the indicated cells stable transduced with shNT or shRNAs against LDHB. b-d, viability assay of shNT- or shLDHB-transduced cells after treated with RSL3 or Erastin, alone or combination with 2 μM Ferrostatin-1(FER1), 40 μM Z-VAD-FMK(ZVF), 20 μM Necrostatin-1 (NEC1), 10 μM Hydroxychloroquine (HCQ). e, immunoblots of the indicated cells transfected with siNT or siLDHB for 48 hours. f-g, flow cytometry of C11 BODIPY mean fluorescence intensity ratio of oxidative channel (FITC 488 nm) versus non-oxidative channel (PE-TEXAS RED 610 nm) in shNT- or shLDHB-transduced cells. A549 cells were treated with 0.5 μM RSL3 or 5 μM Erastin, alone or combined 2 μM FER1 for 6 h and 14 h, while H838 cells with 0.25 μM RSL3 or 2.5 μM Erastin, alone or combined 2 μM FER1 for 6 h and 14 h. h, PTGS2 mRNA levels in A549 and H838 cells after the same treatment as in f-g. i, j, cell death (sytox staining) and lipid peroxidation (liperfluo) assay of the indicated cells transfected for 48 h with siNT or siLDHB and subsequently treated with ML162 in the presence or absence of Liproxstatin-1 (LIP1) for 7 h. k, viability assay of A549 and H838 cells transfected with siNT or siLDHB for 24 h and subsequently transfected with siNT or siSLC7A11/siGPX4 for 48 h. Data are shown as mean ± s.d. (n=3), with the statistical analyses by one-way ANOVA. ns, no significant difference.

Supplementary Figure 5

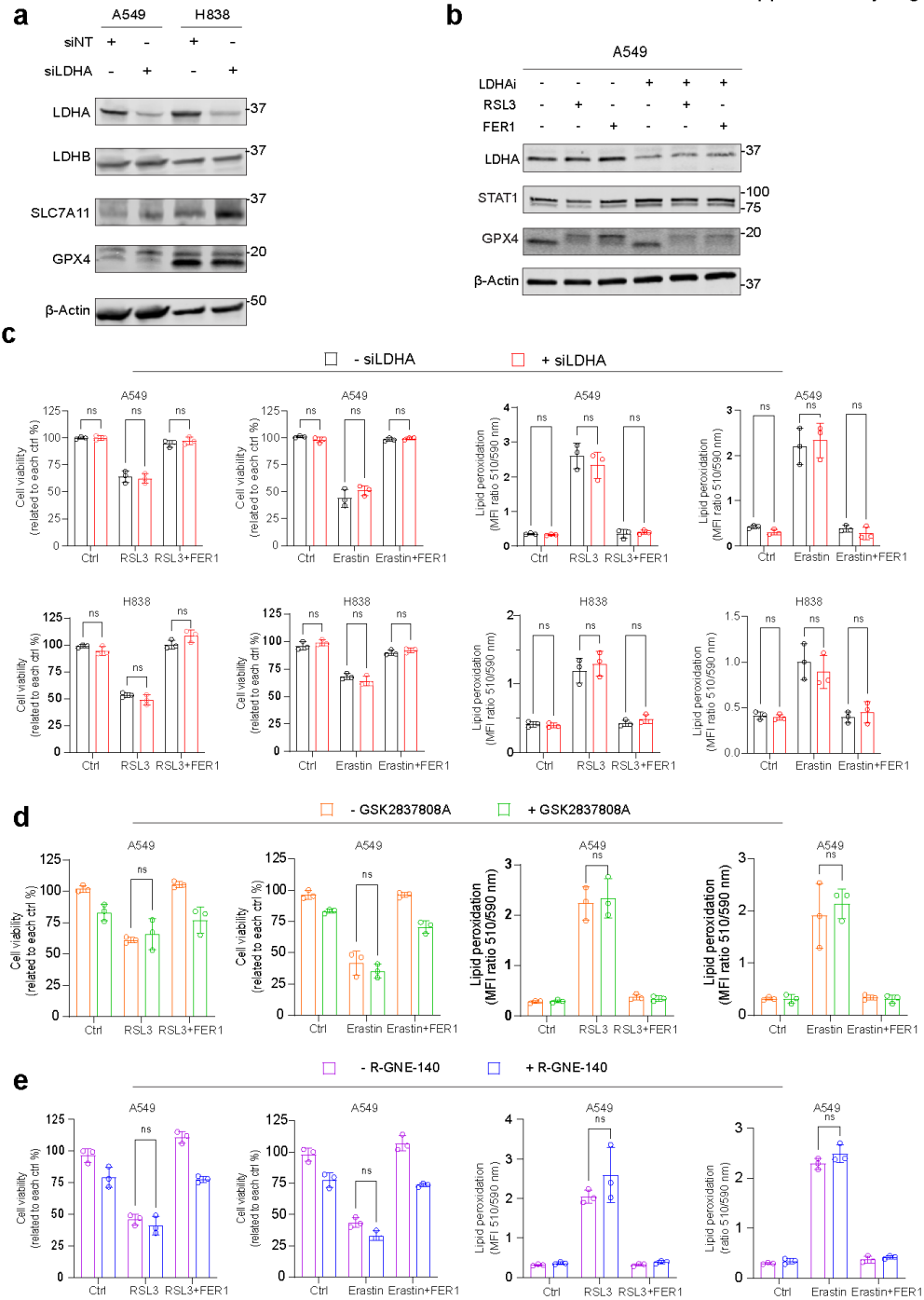

**Figure S5, LDHA is not involved in ferroptosis surveillance in KRAS-dependent lung cancer.** a, immunoblot analysis of the indicated cells transfected with siNT or siLDHA for xx h. b, immunoblot analysis of A549 cells transfected with siNT or siLDHA, followed by treatment with RSL3, alone or combined with FER1. c, viability and lipid peroxidation assay of A549 and H838 cells transfected with siNT or siLDHA and subsequently treated with RSL3, Erastin and FER1,

alone and in combination. Lipid peroxidation levels are shown as the mean fluorescence intensity ratio of the oxidative form versus non-oxidative form. d, viability assay of A549 cells pre-treated with GSK2837808A or vehicle, followed by treatment with RSL3 or Erastin, alone or in combination with FER1. e, viability assay of A549 cells pre-treated with R-GNE-140 or vehicle, followed by treatment with RSL3 or Erastin, alone or in combination with FER1. Data are shown as mean ± s.d. (n=3), with statistical analyses by two-way ANOVA. ns, no significant difference.

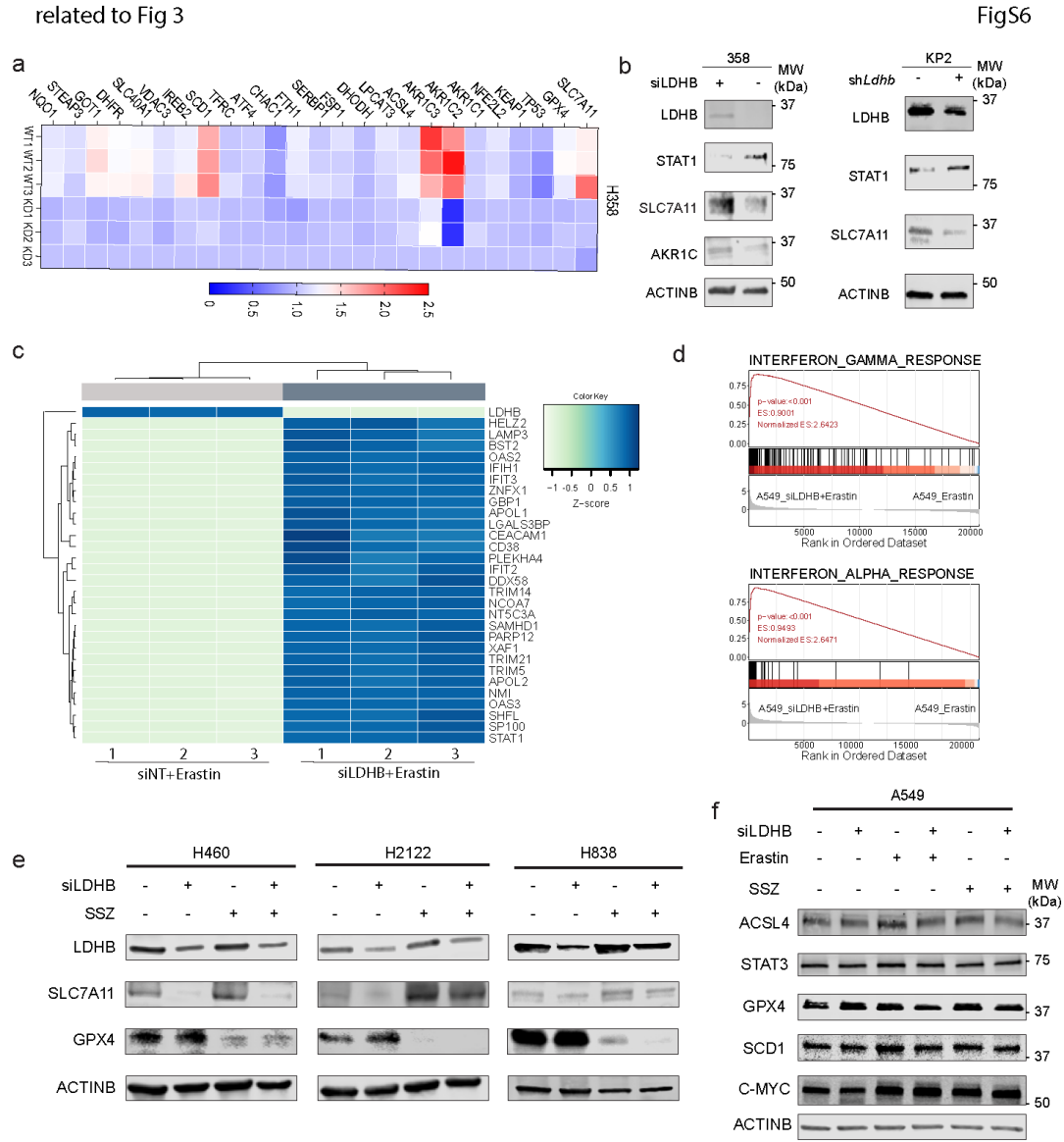

**Figure S6, related to Fig 3.** a, heat map showing the fold change (mRNA levels) of 25 ferroptosis-related genes in siLDHB cells compared to siNT in H358 cells. b, immunoblots of human and mouse LDHB, STAT1, SLC7A11 and AKR1C in the indicated cells. c, the top 30 significantly differentially expressed genes between siNT+Erastin group and siLDHB+Erastin group. Data are presented as z-score values between -1 to 1. d, GSEA plots showing that interferon gamma response and interferon alpha response gene sets are enriched in siLDHB+Erastin cells compared to Erastin cells. e, immunoblots of LDHB, SLC7A11 and GPX4 in H460, H2122 and H838 cells after the indicated treatment. f, immunoblots of ACSL4, STAT3, GPX4, SCD1, and C-MYC in A549 cells after the indicated treatment.

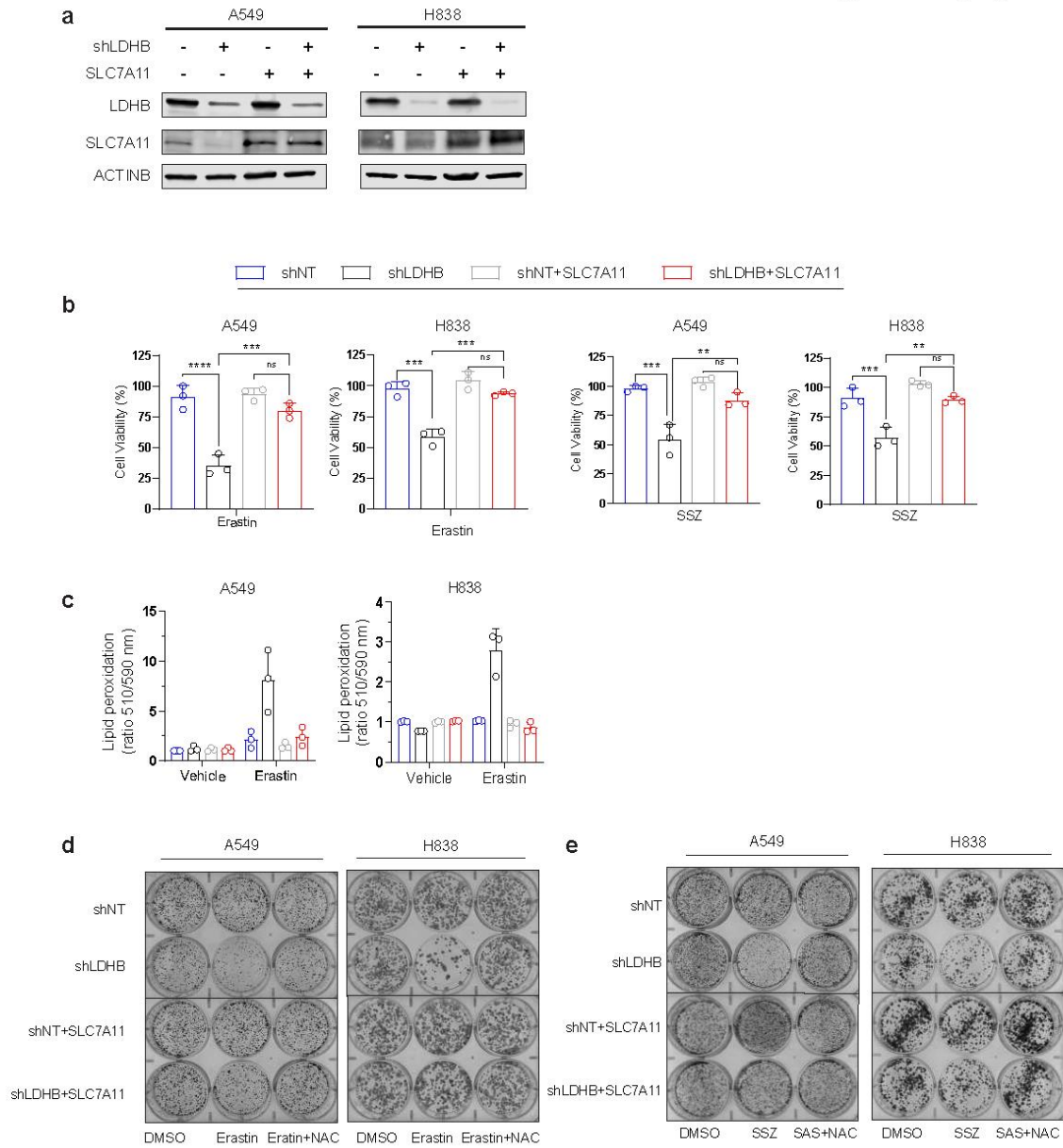

**Figure S7, SLC7A11 overexpression rescues the combinatorial effect of LDHB inhibition plus Erastin or SSZ.** a, immunoblot analysis of A549 expressing shControl or shLDHB) were further transduced with either an empty vector or a SLC7A11-expressing plasmid (pCMV-SLC7A11). b, c, viability and lipid peroxidation assay of A549 and H838 cells treated with Erastin or SSZ for 24 h. Data are shown as mean ± s.d (n=3), with \*P<0.05, \*\*P<0.01, \*\*\*P<0.001 by two-way ANOVA. d-e, clonogenic assay of A549 and H838 cells transduced and treated as indicated.

Supplementary Figure 8

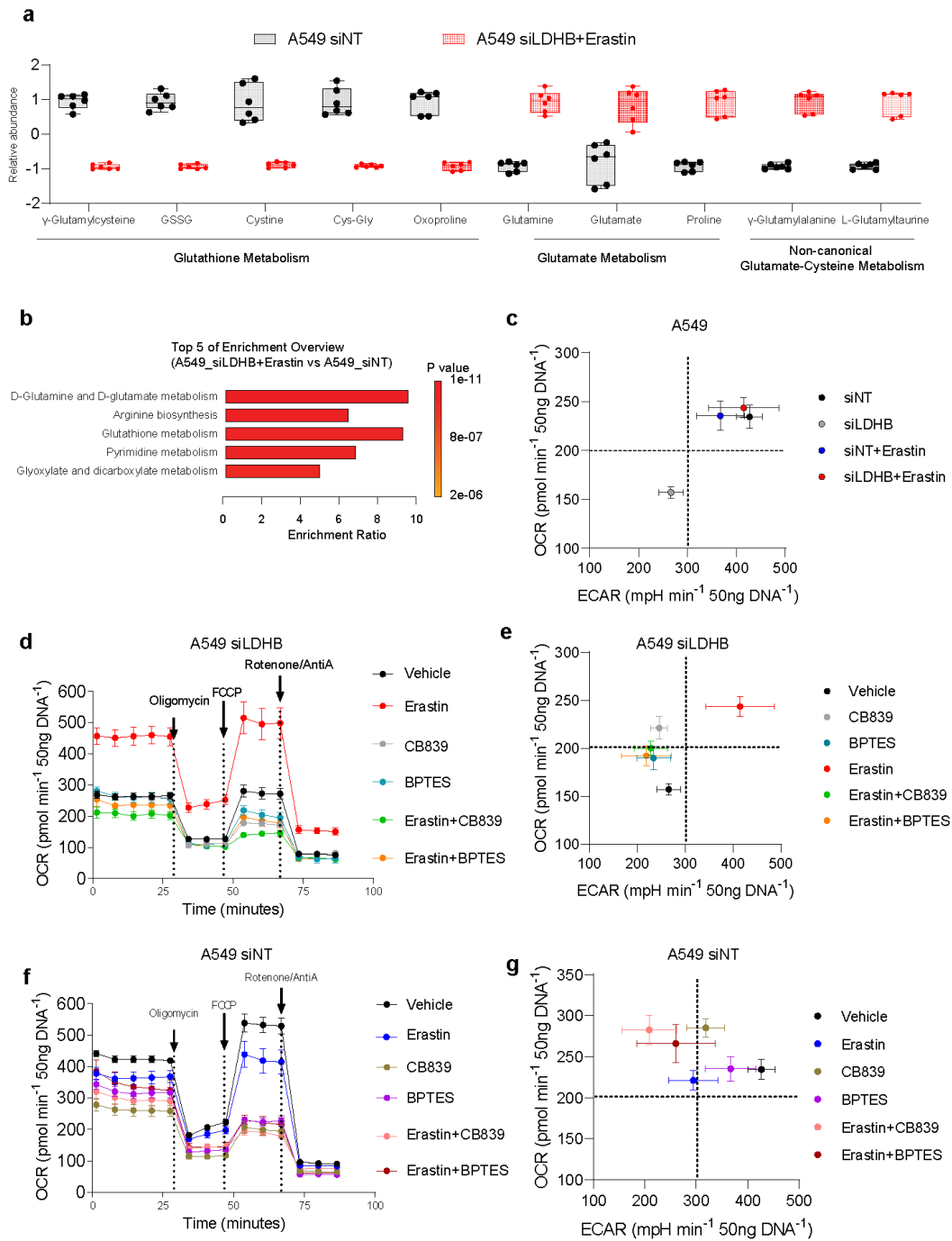

**Figure S8**, related to Figure 4, a, normalized abundance of metabolites. b, the top 5 significantly different metabolism pathways between indicated groups. c, metabolic phenotypes of A549 cells transfected with siNT or siLDHB for 48 h and further treated with Erastin (5 $\mu\text{M}$) for 20 h. d-g, OCR and metabolic phenotypes of A549 cells transfected as in c and further treated for 20 h with Erastin (5 $\mu\text{M}$), CB839 (0.5 $\mu\text{M}$), BPTES (2 $\mu\text{M}$), GPNA (50 $\mu\text{M}$), alone or in combination.

Supplementary Figure 9

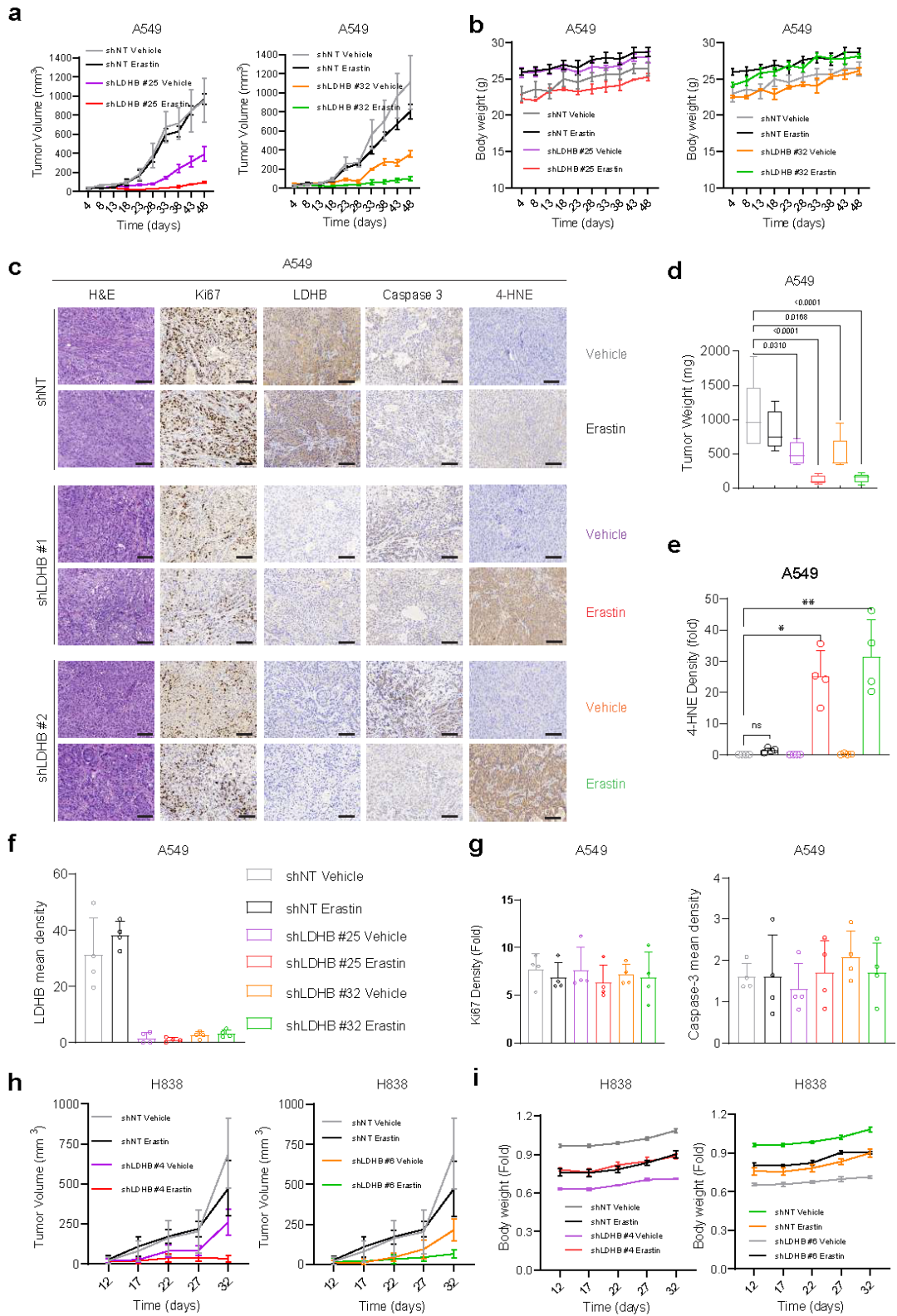

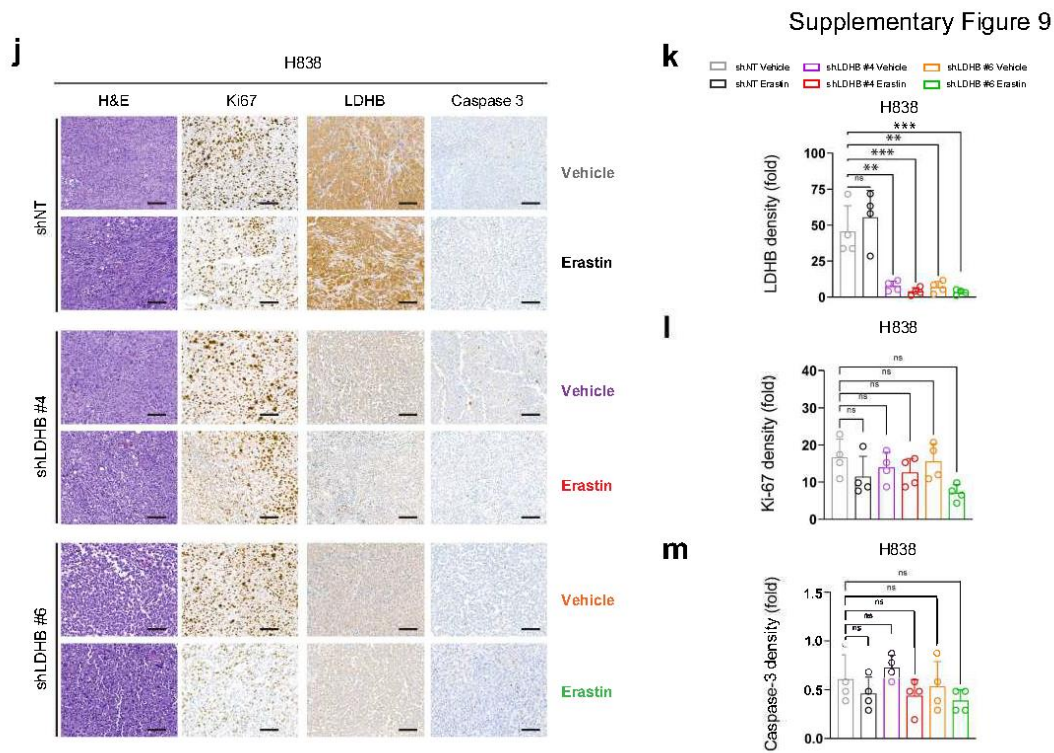

**Figure S9, in vivo efficacy of Erastin in LDHB-deficient KRAS-driven lung cancer models.** a-g, growth curve (a), body weight (b), and IHC analysis (c), and tumor weight (d) of the KRAS-mutant lung cancer A549 stably expressing shNT or shRNA after treated with vehicle, Erastin (30 mg/kg/day). Quantification of IHC staining are shown in e, f, g. h,i growth curve and body weight of the KRAS-mutant lung cancer H838 stably expressing shNT or shRNA after treated with vehicle, Erastin (30 mg/kg/day). k-m, IHC and quantification of H838 xenografts. data are shown as mean ± s.d. of at least three biological repeats (n=3), with \*P<0.05, \*\*P<0.01, \*\*\*P<0.001 by two-way ANOVA. Ns, means no significance.

related to Fig 6

Supplementary Figure 10

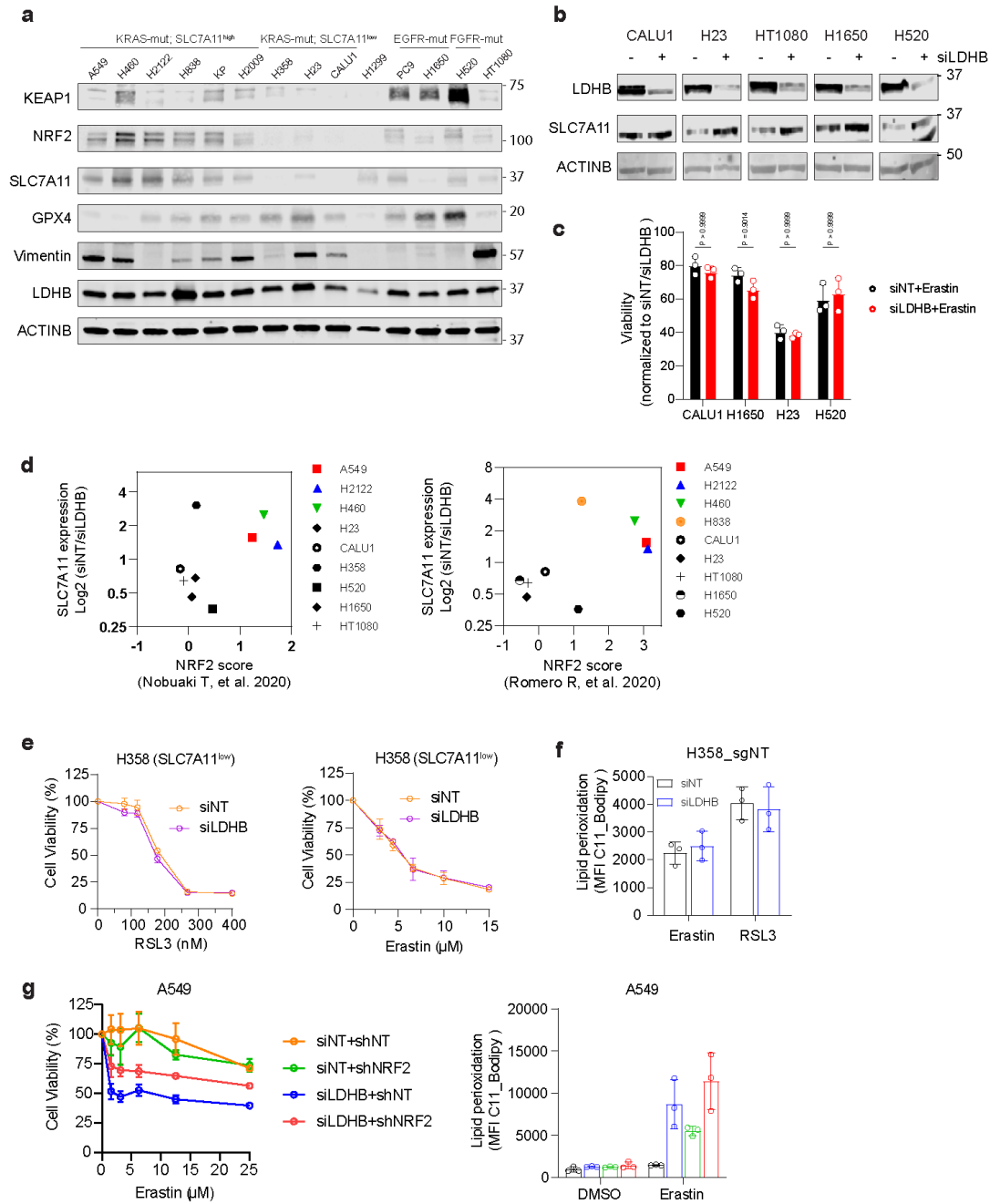

**Figure S10, High levels of SLC7A11 predict the efficacy of LDHB inhibition combined with ferroptosis inducers.** a, immunoblot analysis of KEAP1, NRF2, SLC7A11, and GPX4 in the indicated cell lines. b, immunoblot analysis of LDHB and SLC7A11 in indicated cells transfected with siNT or siLDHB. c, viability assay of CALU1, H1650, H23 and H520 cells transfected with siNT and siLDHB and treated with Erastin (200 nM) for 72 h. d, The SLC7A11 level and the NRF2 score are associated with the sensitivity of cancer cells to combination treatment (LDHBi + Erastin/RSL3/SSZ). The Y-axis represents fold changes of SLC7A11 (siNT/siLDHB). The NRF2 score in the X-axis is based on the indicated studies. e-f, viability assay and lipid peroxidation of H358 cells transfected with either siNT or siLDHB after treated with RSL3 and Erastin for 24 h or 6h (refer to Figure 6c-f), respectively. g, immunoblots of A549 cells stably expressing a doxycycline-inducible (0.5 μM) shControl (tet\_pLKO.1\_shcoo2) or a doxycycline-inducible (0.5 μM) NRF2-specific shRNA (tet\_pLKO.1\_shNRF2#1, shNRF2), followed by further transfection with siNT or siLDHB.

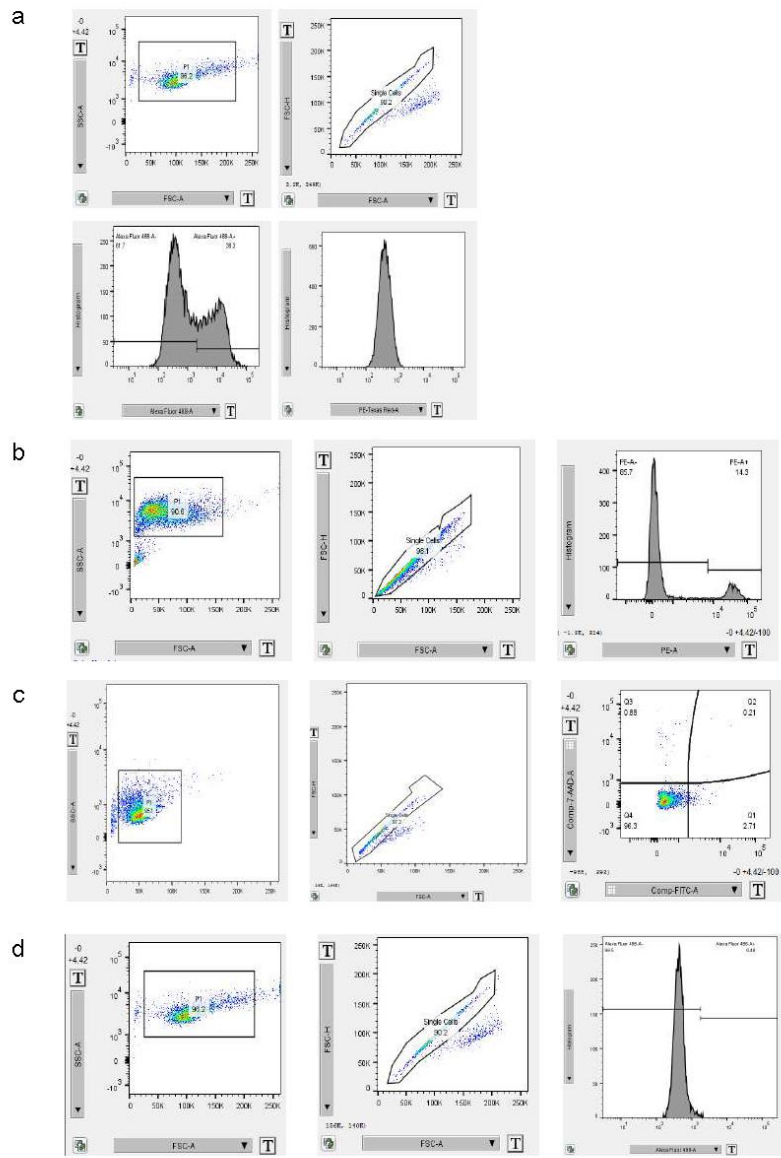

**Figure S11, Gating strategy for Flow Cytometry.** cell population gating was adopted to make sure only single cells were used for analysis. a, C11-BODIPY gating strategy. b, sytox staining of cell death gating strategy. c, Annexin V and PI staining of apoptotic cells gating strategy. d, Liperfluo and DCFDA staining gating strategy.
